# A Systematic Analysis of Mosquito-Microbiome Biosynthetic Gene Clusters Reveals Antimalarial Siderophores that Reduce Mosquito Reproduction Capacity

**DOI:** 10.1101/2020.04.09.034280

**Authors:** Jack G. Ganley, Ashmita Pandey, Kayla Sylvester, Kuan-Yi Lu, Maria Toro-Moreno, Sina Rütschlin, James M. Bradford, Cody J. Champion, Thomas Böttcher, Jiannong Xu, Emily R. Derbyshire

## Abstract

Advances in infectious disease control strategies through genetic manipulation of insect microbiomes have heightened interest in microbially produced small molecules within mosquitoes. Herein, 33 mosquito-associated bacterial genomes were mined and over 700 putative biosynthetic gene clusters (BGCs) were identified, 135 of which belong to known classes of BGCs. After an in-depth analysis of the 135 BGCs, iron-binding siderophores were chosen for further investigation due to their high abundance and well-characterized bioactivities. Through various metabolomic strategies, eight siderophore scaffolds were identified in six strains of mosquito-associated bacteria. Among these, serratiochelin A and pyochelin were found to reduce female *Anopheles gambiae* overall fecundity likely by lowering their blood feeding rate. Serratiochelin A and pyochelin were further found to inhibit the *Plasmodium* parasite asexual blood and liver stages *in vitro*. Our work supplies a bioinformatic resource for future mosquito microbiome studies and highlights an understudied source of bioactive small molecules.

## INTRODUCTION

Over the last decade, efforts to understand host-microbiome interactions mediated by small molecules have significantly increased (Medema, 2018; Milshteyn et al., 2018). Investigation of insect microbiomes has revealed the chemical diversity derived from complex microbial flora within insects (Beemelmanns et al., 2016, 2017, Carr et al., 2012b, 2012a; Ganley et al., 2018; Oh et al., 2009; Reimer et al., 2013). Yet, the majority of natural product studies involving insect-associated microbes focus on a single molecule or group of molecules from individual strains, which are often linked to a particular phenotype or activity (Kroiss et al., 2010; Nollmann et al., 2015; Oh et al., 2009). Bioinformatic tools that predict BGCs is one route to reveal the biosynthetic potential of a microbial community. This approach has been utilized to survey the human gut (Donia et al., 2014) and oral microbiomes (Aleti et al., 2019) as well as plant microbiomes (Helfrich et al., 2018), however it has yet to be applied to insect microbiomes.

Within the insect family there is a pressing need to study *Anopheles* mosquitoes as they include the primary vectors for *Plasmodium* transmission, the causative agent of malaria, as well as other infectious agents. Previous work indicates that the mosquito gut flora is essential for insect development (Coon et al., 2014, 2016), can influence infectious disease transmission (Cirimotich et al., 2011; Ramirez et al., 2014; Stathopoulos et al., 2014), and has the capacity to produce antimalarial small molecules (Ganley et al., 2018; Saraiva et al., 2018). Efforts to control disease transmission through microbial interference of the vector microbiome have proven efficacious for both dengue fever (Jeffery et al., 2009) and malaria (Lovett et al., 2019; Shane et al., 2018; Wang et al., 2017) in laboratory and near-natural environments. Specifically, paratransgenesis studies have illustrated that heterologous expression of antimalarial (Shane et al., 2018; Wang et al., 2017) or insecticidal (Lovett et al., 2019) proteins or peptides in microbiome species can efficiently reduce parasite load or influence mosquito survival, respectively. Thus far, these approaches have primarily employed insect venom proteins as the inhibitory agent, but antimalarial or insecticidal small molecules are also attractive candidates (Kajla, 2019).

In this study, we completed an extensive bioinformatic analysis of the small molecule BGCs from mosquito-associated bacteria. We mined through 33 mosquito-associated bacterial genomes and identified over 700 putative BGCs. Further bioinformatic analysis indicated that siderophores, small molecules secreted to sequester and replenish intracellular iron stocks, were highly represented within this bacterial sample set. Siderophores have been reported with diverse bioactivities (Johnstone and Nolan, 2015) including *in vitro* and *in vivo* anti-*Plasmodium* activity (Atkinson et al., 1991; Fritsch et al., 1985; Shanzer et al., 1991). In the mosquito microbiome, we identified eight different siderophore scaffolds through a mass spectrometry analysis and subsequently obtained six purified compounds for bioactivity studies. The siderophores serratiochelin A (Araz and Budzikiewicz, 1994) and pyochelin (Cox et al., 1981) exhibited adverse effects on the blood feeding propensity and overall fecundity of female *A. gambiae* mosquitoes. Additionally, these compounds were identified as inhibitors of the liver- and asexual blood-stages of *Plasmodium* infection. Together, we find that bacterial siderophores are important mosquito-microbiome molecules and highlight two siderophores with activities against the mosquito vector and *Plasmodium* parasite. Identifying metabolites with these properties within the mosquito-microbiome lays the foundation for future paratransgenesis campaigns to reduce disease transmission and mosquito competence.

## RESULTS AND DISCUSSION

### *In silico* identification of biosynthetic gene clusters from mosquito gut microbiome

We compiled publicly available bacterial genomes that were originally isolated from mosquitoes for analysis. This afforded 29 bacterial strains from *Anopheles* mosquitoes (*A. gambiae, A. stephensi, A. arabiensis*, and *A. sinensis*) and 4 strains from *Aedes* mosquitoes (*Ae. aegypti* and *Ae. albopictus*) (**Data Set S1A**). *Aedes* mosquitoes transmit avian malaria, *P. gallinaceum* (Alavi et al., 2003), as well as human infectious agents including dengue virus, Zika virus, and others (2016). As a relevant disease vector, *Aedes*-associated bacteria were also included in our data set. Due to the limited amount of genome sequencing data on mosquito-associated bacteria, especially when compared to human microbiomes, our sample size is smaller than analogous studies (Aleti et al., 2019; Donia et al., 2014; Helfrich et al., 2018), however this enabled an in-depth bioinformatic analysis. Despite this size, our data set well-represents the unique gut microbiome of mosquitoes. Our previous metagenomic study demonstrated that within the gut microbiota of field-caught *A. gambiae* mosquitoes four days post blood-meal, 5 bacterial genera (*Serratia, Elizabethkingia, Acinetobacter, Enterobacter*, and *Pseudomonas*) constitute 84 % of the 16S rRNA reads (Wang et al., 2011), all of which are represented within our sample set.

A phylogenetic tree comparing the 33 bacterial strains was generated (**Figure 1**). The best-represented bacterial phylum in our sample set was *Proteobacteria* (16 γ-*Proteobacteria*, 1 β*-Proteobacteria*, and 3 α*-Proteobacteria*), consistent with reports showing *Proteobacteria* are often the overwhelming microbial phylum in the midgut of field-caught *Anopheles* mosquitoes (Wang et al., 2011). Species in the *Bacteroidetes* phylum, which include *Elizabethkingia* spp., were the second most recurrent group (4 *Elizabethkingia* spp. and 1 *Sphingobacterium* sp.) in our analysis. *Elizabethkingia* spp. are well-established mosquito symbionts that are the dominant species within midguts 4- and 7-days post-blood meal (PBM) as well as 7-days after sugar feeding of *A. gambiae* mosquitoes (Wang et al., 2011).

**Figure 1.**
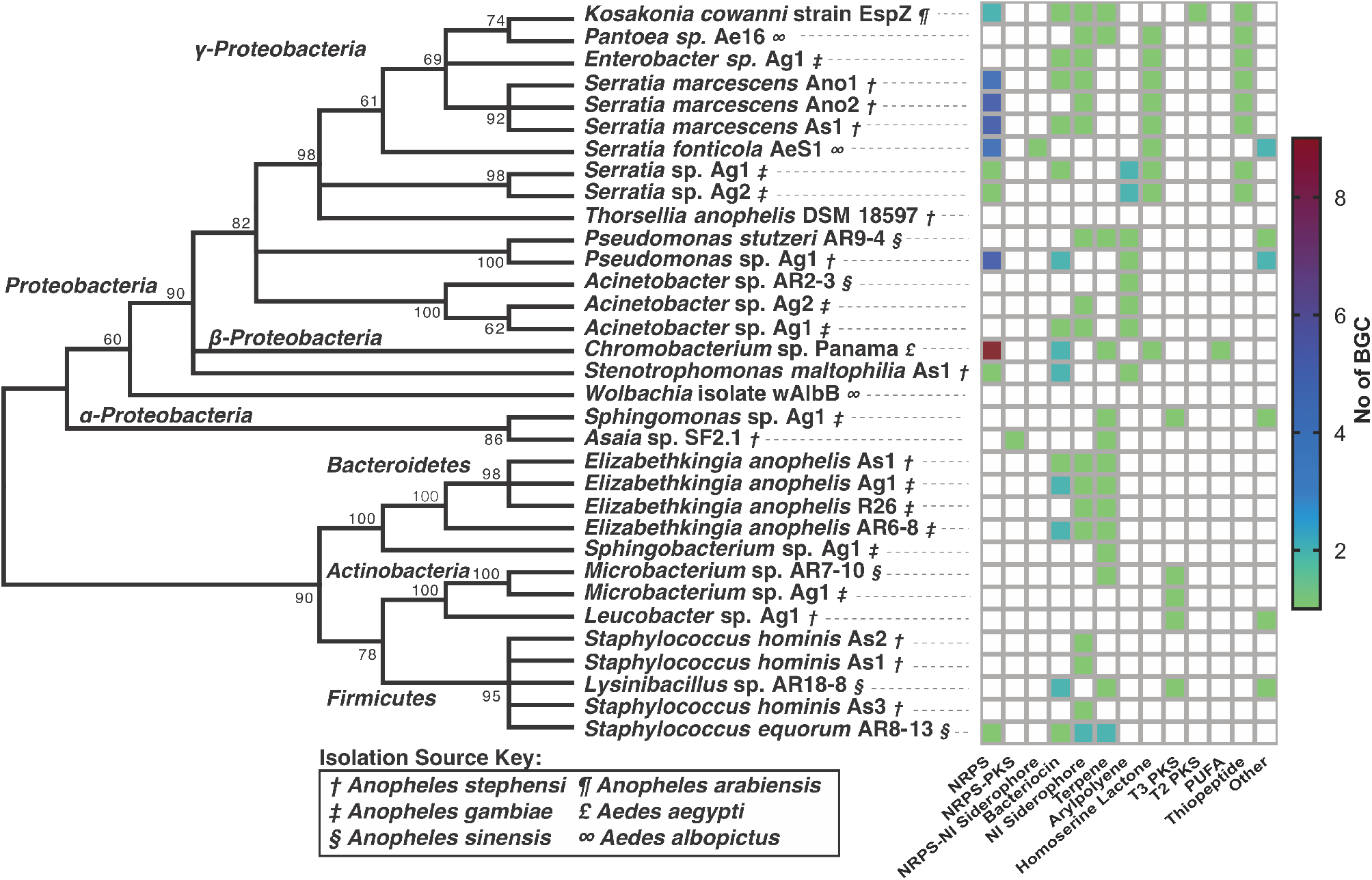
Summary of the Mosquito-Microbiome Strains Examined and their Biosynthetic Potentials. Maximum likelihood phylogenetic tree of 16S rRNA of all 33 strains examined in this study and their biosynthetic potential. The heat map to the right of the phylogenetic tree indicates the number of each type of BGC identified by antiSMASH for each strain. White boxes indicate zero BGCs from that class were found within the genome. Symbols next to each strain name indicate the original mosquito isolation source (Isolation Source Key).

Each bacterial genome was subjected to a bioinformatic analysis to predict their biosynthetic potentials. ClusterFinder (Cimermancic et al., 2014), a Markov model-based probabilistic algorithm that identifies BGCs from known and unknown classes, was used to survey the genomes. From our analysis, 719 total BGCs were identified by ClusterFinder. Of the 719 BGCs, antiSMASH (Blin et al., 2017), a bioinformatic tool to detect BGCs in defined classes, identified 135 BGCs of various types (**Figure 2A**, **Data Set S1B**). The heat map in **Figure 1** indicates the classes of antiSMASH BGCs as well as the number of BGCs in each strain. We observed that nonribosomal peptide synthetase (NRPS)-independent (NI) siderophores, terpenes, and bacteriocins were well-distributed phylogenetically, while classes like aryl polyenes and homoserine lactones are limited to distinct phyla. The most frequent BGCs were NRPS clusters (38/135), which were almost exclusively found in *Proteobacteria*, specifically *Chromobacterium*, *Serratia*, and *Pseudomonas* spp. Bacteriocins (20/135), NI siderophores (18/135), and terpenes (15/135) were also prevalent BGCs (**Figure 2B**). When present within genomes, NRPS clusters were found in higher numbers (>4) when compared to other classes. This abundance hints at potentially important roles for molecules synthesized by NRSPs within these organisms. Conversely, there was a surprising lack of modular type-I polyketide synthases (PKSs) detected in this analysis, a class often reported in human and plant microbiome studies (Donia et al., 2014; Helfrich et al., 2018). Specifically, only two modular type-1 PKS BGCs (**Data Set S2**, Clusters Asaia15 and Chromobacterium11) were identified.

**Figure 2.**
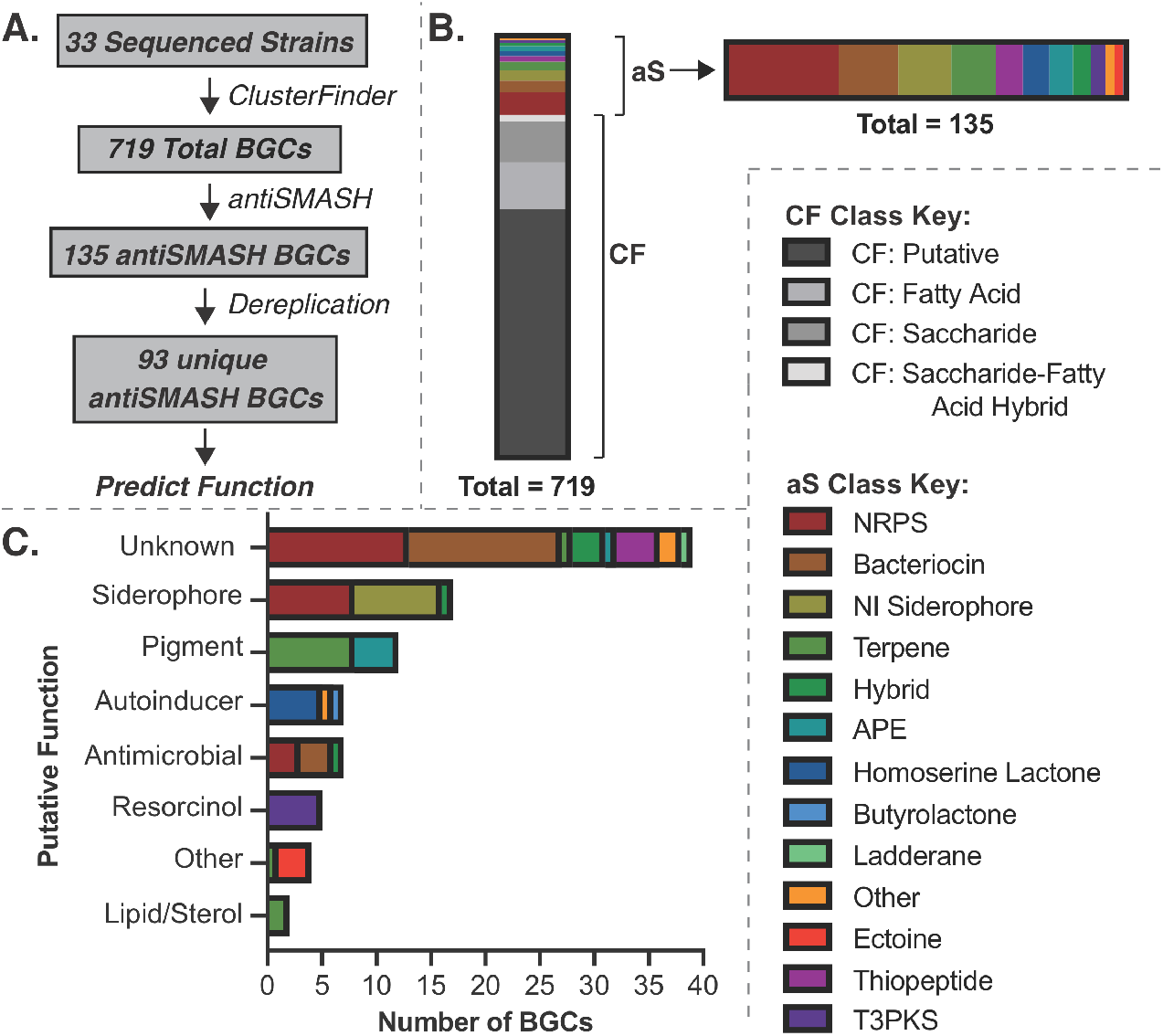
A Summary of the BGCs from Mosquito-Microbiomes and Predicted Bioactivities. (**A**) Overview of the bioinformatic analysis. (**B**) Summation of all BGCs indicating the amounts of antiSMASH (aS) and ClusterFinder (CF) BGCs and the type of each cluster with colors distinguishing the different classification. (**C**) Bar chart indicating the predicted function of the 93 unique antiSMASH clusters with colors distinguishing the antiSMASH classification.

To evaluate the associated secondary metabolites from BGCs we assigned putative physiological functions to the 135 antiSMASH clusters. Since BGCs within the same antiSMASH classification can encode for various types of metabolites with diverse bioactivities, we compared each cluster to the phylogenetically-closest characterized natural product cluster with established biological activity (**Data Set S1C**, see **Materials & Methods** for more details). Natural products can have multiple physiological functions, including functions that have yet to be discovered, therefore we binned by the best-characterized function. This approach was especially useful for identifying redundant BGCs, as some of the strains within our dataset contain nearly identical genomes (i.e. *Serratia* sp. Ag1 and *Serratia* sp. Ag2, 98.5523% symmetrical identity). After excluding the redundant BGCs (**Data Set S1D**), the 93 remaining unique antiSMASH BGCs were ascribed to one of the following predicted functions: Unknown, siderophore, pigment, autoinducer, antimicrobial, other, resorcinol, and lipid/sterol (**Figure 2C**). From this analysis, it was apparent that the sample set contains many BGCs that likely produce uncharacterized molecules, as more than one third of the BGCs have no closely associated characterized BGC. The antiSMASH clusters with unknown predicted function encompassed various classes, including NRPS, hybrid, terpene, bacteriocin, and thiopeptide BGCs. This suggests that these strains are untapped sources of undiscovered molecules. In particular, *Chromobacterium* sp. Panama (Ramirez et al., 2014), *Serratia fonticola* AeS1, *Kosakonia cowanii* Esp_Z (Cirimotich et al., 2011), and *Lysinibacillus* sp. AR18-8 have a disproportionately high amount of unknown BGCs. Together these strains contain approximately half of the unknown BGCs (19/39), making them attractive candidates for future natural product discovery campaigns. To further validate our approach, we complemented our bioinformatics analysis with a mass spectrometry study.

### Identification of mosquito-gut microbiome siderophores

Siderophores were the most abundant functional prediction from our analysis, constituting 17 unique BGCs, 7 of which were detected in more than one strain. Canonically, siderophores are synthesized and secreted by organisms in low-iron environments to sequester and retrieve Fe^3+^ for typical cellular processes like respiration and DNA synthesis (Lankford and Byers, 1973). This function is particularly compelling in the context of mosquito biology as the iron-binding small molecules would directly interact with the *Plasmodium* parasite in the midgut during the extracellular sexual reproduction stage. To investigate these possible siderophores, six mosquito-microbiome strains (covering 9 of the 17 siderophore BGCs) were acquired and grown in standard versus iron-limited media. Since FeCl_3_ typically arrests siderophore production, an Fe-dependent change in a metabolite level can facilitate siderophore detection by mass spectrometry. Analyzed strains included *Serratia* sp. Ag2, *Serratia* sp., *Enterobacter* sp. Ag1, *Pseudomonas* sp. Ag1, *Acinetobacter* sp. Ag1, and *E. anophelis* Ag1 (**Data Set S1E**). *Serratia* sp. (Ganley et al., 2018) was used in place of *S. marcescens* AS1 as it has 99 % 16S rRNA identity and contains identical siderophore BGCs. Differential metabolomics was then employed to identify metabolites upregulated in iron-deficient conditions, which were further investigated by tandem mass spectrometry (MS/MS) analysis (**Figure 3, Fig. S1, Data Set S3**). This approach identified serratiochelin A and B from *Serratia* sp. (**Figure 3A**), pyochelin (Ganley et al., 2020) and the related metabolite dihyroaeruginoic acid from *Pseudomonas* sp. Ag1 (**Figure 3B**), and aerobactin from *Enterobacter* sp. Ag1 (**Figure 3C**). As predicted by our BGC analysis, some strains produce multiple siderophores. Additional targeted metabolomic searches identified five other siderophore scaffolds including, chrysobactins (*Serratia* sp., **Figure S2**), enterobactin (*Serratia* sp. Ag2, **Figure S3A–C**), acinetoferrin (*Acinetobacter* sp. Ag1, **Figure S1F**), bisucaberin (*E. anophelis* Ag1, **Figure S3D–G**), and pyoverdine (*Pseudomonas* sp. Ag1, **Figure S4**) (**Figure 3D**). Interestingly, *Pseudomonas* sp. Ag1 produces a previously undiscovered pyoverdine named pyoverdine Ag1 in addition to pyochelin. We predict a putative pyoverdine Ag1 structure by combining MS/MS and biosynthetic predictions (**Figure S4**), however full structure elucidation awaits 2D NMR characterization.

**Figure 3.**
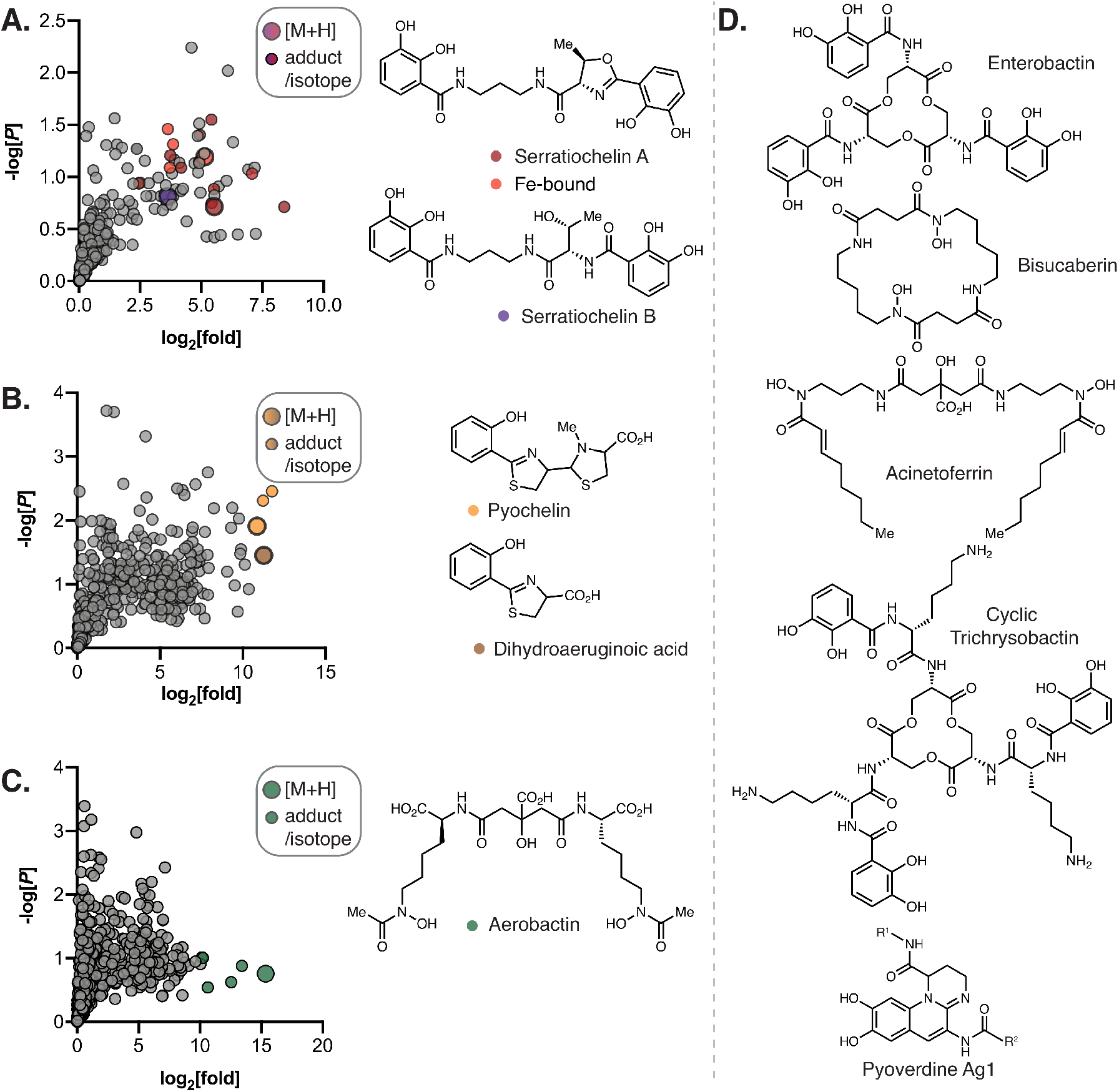
Overview of the Siderophores Identified in this Study, Including Differential Metabolomics Results from Three Bacterial Strains Analyzed. (**A**) Volcano plot of upregulated ions of *Serratia* sp. grown in iron-deficient media. (**B**) Volcano plot of upregulated ions of *Pseudomonas* sp. Ag1 grown in iron-deficient media. (**C**) Volcano plot of upregulated ions of *Enterobacter* sp. Ag1 grown in iron-deficient media. (**D**) Structures of other siderophores identified in this study. All stereochemistry is predicted based on previously published structures and biosynthetic predictions.

Our study provides a blueprint of siderophores employed by mosquito-associated bacteria. While some of these strains are well-studied for their chemical potential, few reports exist on the secondary metabolites of the emerging nosocomial pathogen *E. anophelis*. We found that this bacterium contains a BGC homologous to a desferrioxamine-like BGC that is conserved across species. Additionally, the metabolomic and MS/MS results indicated that the cyclic hydroxamate siderophore bisucaberin is produced in metal-depleted *E. anophelis* Ag1 culture (**Figure S3D–H**). Bisucaberin was initially described in marine species of *Aliivibrio* (Winkelmann et al., 2002) and *Alteromonas* (Takahashi et al., 1987). In *Shewanella algae*, bisucaberin is simultaneously produced with avaroferrin and putrebactin via the same BGC and ratios of these compounds were controlled by substrate availability (Böttcher and Clardy, 2014; Rütschlin et al., 2017). Interestingly, entomopathogenic *Xenorhabdus szentirmaii* only produced putrebactin and avaroferrin (Hirschmann et al., 2017), while in our analysis of *E. anophelis* Ag1 only bisucaberin was detected, suggesting great diversity and fine control of the ratio of these siderophores by different species. Overall, all of the siderophores identified by MS were putatively assigned to a BGC (**Data Set S1E**) and then prioritized for bioactivity studies. Among the siderophores identified, six compounds were attained through either isolation from the producing bacterial strain, *in vitro* production and isolation, or purchasing, (see **Resources Table** for siderophore source).

### The mosquito-microbiome siderophores, serratiochelin A and pyochelin reduce blood feeding propensity and overall fecundity

Serratiochelin A, pyochelin, and aerobactin could be produced or purchased in quantities sufficient for bioactivity assays in *A. gambiae*, including evaluation of mosquito survival, feeding rate, and fecundity. In these experiments, siderophores (100 μM) were provided to 100 female mosquitoes with a sugar-meal for three days before a blood meal. Engorged female mosquitoes were then separated and dissected three days thereafter (**Figure 4A**). Over the three days of siderophore supplementation, survival rates of female mosquitoes were monitored. After averaging survival of siderophore-fed female mosquitoes across three biological replicates, we observed no consistent reduction in survival among siderophore-fed female mosquitoes (**Figure 4B**), however serratiochelin A and pyochelin showed some toxicity in survival curves of individual replicates (**Figure S5**). Additionally, we recorded the percentage of female mosquitoes that took a blood meal post-siderophore feeding. Compared to the DMSO vehicle control where 53% of mosquitoes fed, pyochelin caused a significant reduction in blood feeding propensity across the three biological replicates, where only 37 % took blood meals (**Figure 4C**). Since the three biological replicates had variable sample sizes, we pooled the blood feeding propensity to observe the total percent of engorged females (**Figure 4D**). After pooling the data across the three biological replicates, serratiochelin A and pyochelin both caused a significant reduction in blood feeding propensity at the population level, suggesting the general health of these mosquitoes may be affected by the compounds. However, survival of the engorged females was monitored post-blood meal from days 4–6 and no significant reduction in health was observed (**Figure 4E**). Next, the number of eggs in each mosquito ovary was assessed after dissection of the engorged females. Compared to the DMSO vehicle control, there was no significant change in the average number of eggs per mosquito for any of the siderophores across the three biological replicates (**Figure 4F**). Although the average number of eggs per mosquito had no significant reduction among blood fed mosquitoes, serratiochelin A and pyochelin both caused greater than a 2-fold decrease in the population’s overall fecundity (total egg number/total starting mosquitoes) when compared to the DMSO vehicle (**Figure 4G**). Since blood meals are required for egg production and pathogen transmission, the observed reduction in blood feeding propensity is likely the main contributors to the overall fecundity reduction. This effect may influence the transmission capacity of mosquito-borne infectious diseases, like malaria. This strategy was recently highlighted by Vosshall and colleagues. Specifically, they discovered compounds that target mosquito peptide receptors and suppress attraction and blood-feeding on live hosts (Duvall et al., 2019). This further illustrates the appeal of the blood-feeding suppression demonstrated by both serratiochelin A and pyochelin.

**Figure 4.**
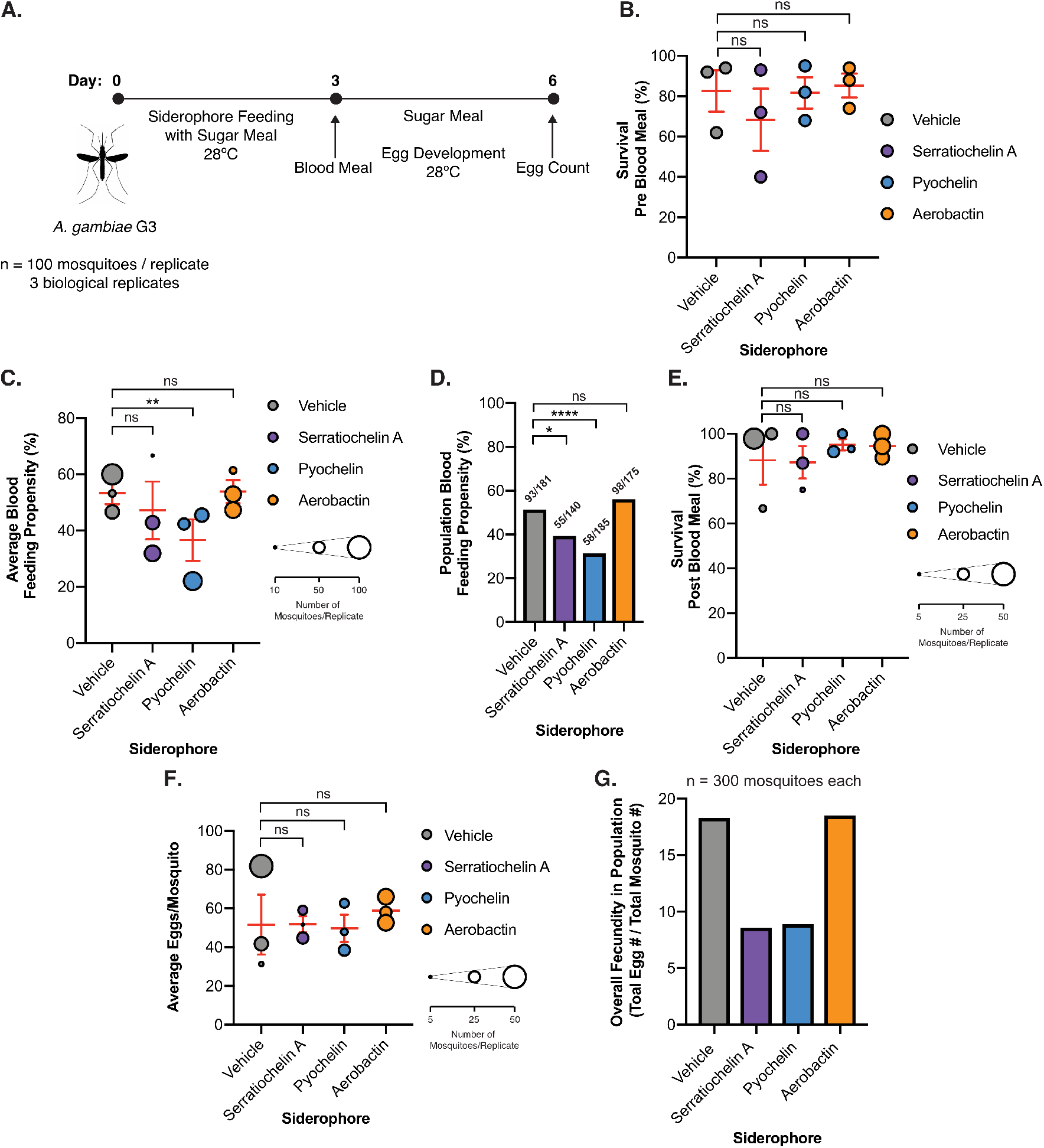
Evaluation of Mosquito-Microbiome Siderophore Activity Against *A. gambiae* Survival, Blood Feeding, and Fecundity. (**A**) Overview of mosquito survival and fecundity assays. For each experiment, 100 female mosquitoes were used, and three biological replicates were completed. (**B**) Average survival of female mosquitoes (at t = 3 days) supplemented with siderophores (100 μM) pre-blood meal. Each dot represents a biological replicate (n = 100). The red bars represent the mean +/− SEM of the three biological replicates (Two-way ANOVA multiple comparisons analysis, ns = not significant). (**C**) Average blood feeding propensity for female mosquitoes post-siderophore feeding. Each dot represents a biological replicate and the size of each dot represents the number of mosquitoes exposed to a blood meal. The red bars represent the mean +/− SEM of the three biological replicates (Two-way ANOVA multiple comparisons analysis, ** *P* < 0.01). (**D**) Total blood feeding propensity at the population level. Each bar represents pooled data for the three biological replicates. The fractions above each bar represent (mosquitoes fed/mosquitoes exposed to a blood meal), (unpaired t-test, * *P* < 0.05, **** *P* < 0.0001). (**E**) Average survival of female mosquitoes (from t = 4–6 days) post-blood meal. Each dot represents a biological replicate and the size of each dot represents the number of mosquitoes exposed that took a blood meal (Two-way ANOVA multiple comparisons analysis). (**F**) Average number of eggs laid per blood-fed female mosquito. Each dot represents a biological replicate and the size of each dot represents the number of blood fed mosquitoes analyzed (Two-way ANOVA, multiple comparisons analysis). (**G**) Overall fecundity of female mosquitoes at the population level when supplemented with various siderophores for 3 days. In total, 300 mosquitoes (3 biological replicates at 100 female mosquitoes each) were supplemented with a vehicle control or a siderophore. Serratiochelin A and pyochelin reduced the overall fecundity by over 50% due to a combination of toxicity and lowered blood feeding propensity.

### Serratiochelin A and pyochelin inhibit *Plasmodium* parasites

All six of the isolated or purchased siderophores were screened for activity in standard anti-*Plasmodium* assays that are available in high-throughput format. As an initial screen, siderophores (10 μM) were tested for inhibition of *P. berghei* (rodent-infective) parasite load in HuH7 hepatoma cells (**Figure 5A**). Three siderophores that significantly reduced parasite load in the initial screen were selected for dose-response analysis. Serratiochelin A and pyochelin inhibited *P. berghei* parasite load with EC_50_ values of 1.6 μM and 510 nM, respectively (**Figure 5B**), while no concentration dependence of inhibition was observed for aerobactin (**Figure S6D–E**). Importantly, serratiochelin A and pyochelin exhibit no HuH7 cytotoxicity under our assay conditions (**Figure S6A–C**). Evaluation of cytotoxicity is a necessary control to establish that *P. berghei* inhibition is not due to host cell toxicity. We further found that serratiochelin A and pyochelin inhibit the blood-stage of *P. falciparum* (human-infective) with EC_50_ values of 10 and 6.6 μM, respectively (**Figure 6C**). Together, this demonstrates inhibition of multiple species and stages of the *Plasmodium* life cycle. Activity against *Plasmodium* gametocyte or mosquito stages is an especially attractive activity for candidate small molecules to reduce malaria transmission. Importantly, iron chelators including FBS0701 and deferoxamine, have been shown to inhibit gametocyte development (Ferrer et al., 2015). This suggests that other compounds that reduce iron levels, like siderophores, have potential to block the transmission of malaria.

**Figure 5.**
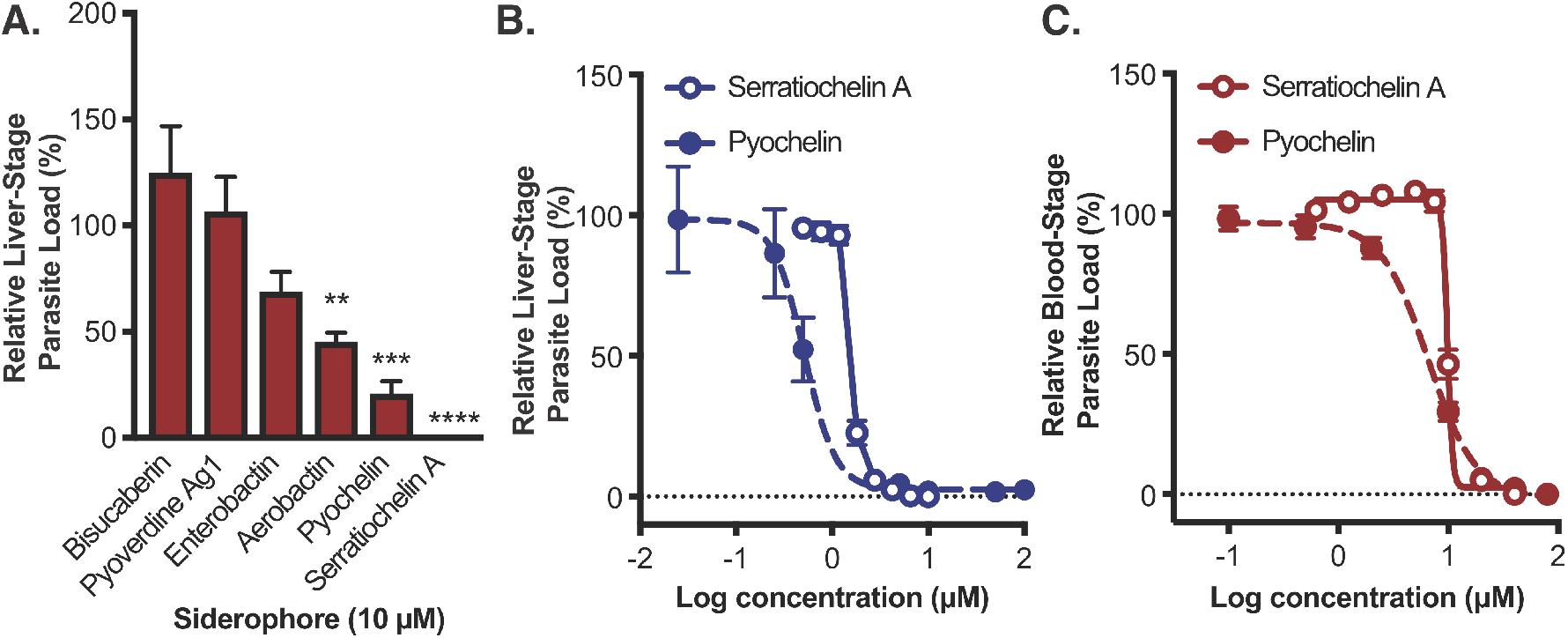
*In vitro* Evaluation of Mosquito-Microbiome Siderophores for Anti-*Plasmodium* Activity. (**A**) Activity of siderophores at 10 μM against *P. berghei* ANKA infection of human HuH7 hepatocytes (One-way ANOVA, Dunnett’s multiple test comparison, ** *P* < 0.01, *** *P* < 0.001, **** *P* < 0.0001). (**B**) Inhibition of *P. berghei* ANKA parasite load in human HuH7 hepatocytes by serratiochelin A (solid line, open circles) and pyochelin (dotted line, filled circles), with EC_50_ values of 1.6 μM and 510 nM, respectively. (**C**) Inhibition of *P. falciparum* 3D7 parasite load in human red blood cells by serratiochelin A (solid line, open circles) and pyochelin (dotted line, filled circles), with EC_50_ values of 10 and 6.6 μM, respectively.

To evaluate whether the siderophores included in our study are produced within mosquitoes, lab-reared *A. gambiae* mosquitoes were analyzed to determine endogenous serratiochelin A and pyochelin levels. Neither sugar-fed male/female mosquitoes nor blood-fed female mosquitoes contained serratiochelin A or pyochelin at detectable concentrations (<10 pg per mosquito) despite containing known producing strains. However, we detected ~ 0.7 ng of serratiochelin A per mosquito in lab-reared female *A. stephensi* mosquitoes (**Figure S7**), indicating that this siderophore is capable of being produced within mosquitoes. Thus, from our initial survey of over 700 mosquito microbiome BGCs, we discovered numerous microbial metabolites including a natural product with activity against multiple *Plasmodium* life cycle stages that is additionally produced within live mosquitoes. The adverse effects of serratiochelin A and pyochelin against *A. gambiae* survival, blood-feeding rates, and overall fecundity, as well as the potential for anti-*Plasmodium* activity, makes these small molecules possible candidates for mosquito population control and transmission control of the *Plasmodium* parasite.

### Potential value of siderophores in a future paratransgenesis campaign

Current paratransgenesis techniques involve the heterologous expression of one or multiple exogenous biomacromolecules that reduce parasite load within *Anopheles* mosquitoes (Shane et al., 2018; Wang et al., 2017). Previous studies have employed *S. marcescens* AS1 as a heterologous host due to its natural occurrence within the *A. stephensi* microbiome and because of its ability to vertically disperse through multiple generations of mosquitoes in a laboratory setting. Expression of various antimalarial effector molecules in *S. marcescens* As1 dramatically reduces parasite load within *A. stephensi* making this an exciting avenue for possible malaria control (Wang et al., 2017). Siderophores are also attractive candidates for this campaign as their well-studied BGCs are relatively short in base-pair length in comparison to large lipopeptide and macrocycle BGCs thus, facilitating efforts to heterologously express and/or manipulate siderophore BGCs for paratransgenesis applications. The combined reduction in overall fecundity with the anti-*Plasmodium* activity of serratiochelin A enhances its future potential as a paratransgenesis tool. Fortuitously, wild type *S. marcescens* AS1 naturally contains the BGC to produce serratiochelin A and therefore, genetic manipulation to constitutively express this BGC in *S. marcescens* AS1 is a feasible alternative or additive approach to current efforts.

Gaining a critical understanding of the repertoire of small molecules within ecological niches is a principal feat when attempting to understand or manipulate the chemical dynamics of a microenvironment. Within the gut of mosquitoes, a complex matrix of inter- and intra-kingdom interactions exist. Advances in bioinformatics, metagenomics, metatranscriptomics, metabolomics, bacterial isolation techniques, and many other areas has facilitated our ability to identify small molecules produced within microbiomes and the BGCs enabling their production. The ability to resolve and manipulate this ecosystem has promising implications for reducing the transmission capacity of various infectious diseases.

The work described herein provides a detailed bioinformatic analysis of the BGCs of prominent mosquito-associated bacterial species and serves as a useful resource for future studies exploring various aspects of chemical ecology within mosquitoes. Our efforts focused on a fraction of the numerous groups of small molecules that are potentially produced within mosquitoes. In addition to highlighting the potential to discover novel small molecules within this system, we identified two anti-*Plasmodial* compounds that hinder the overall reproduction capacity of *Anopheles* mosquitoes. Taken together, this work encourages future endeavors to understand and potentially manipulate the chemical ecology within mosquitoes.

## Supporting information

Supplementary Data Set 1

Supplementary Data Set 2

Supplementary Data Set 3

## AUTHOR CONTRIBUTIONS

J.G.G. performed bioinformatic analyses, siderophore isolations and identification, and mass spectroscopy studies, A.P. and C.J.C., performed all mosquito assays., K.S. and M.T-M. performed liver-stage and cytotoxicity studies; S.R. and T.B. prepared chemical authentic standards and provided siderophore expertise; K.L. performed blood-stage assays; J.M.B. performed isolation and characterization of siderophores; J.G.G., J.X., and E.R.D designed the experiments and analyzed the data; J.G.G. and E.R.D wrote the manuscript; and all authors commented on the manuscript.

## ACKNOWLEDGMENTS

We thank the American Association for the Advancement of Science (Marion Milligan Mason awarded to E.R.D) and Duke University for supporting this research. J.G.G. is grateful for support from a Burroughs-Wellcome fellowship from the Duke University Chemistry Department. J.X was supported by the National Institutes of Health (SC1AI112786) and the National Science Foundation (#1633330). We thank Dr. Peter Silinski for mass spectrometry assistance and Dr. David Gooden for preparative HPLC use. The following reagents were obtained through BEI Resources Repository, NIAID, NIH: *Plasmodium falciparum*, Strain 3D7, MRA-102, contributed by Daniel J. Carucci; *Leucobacter* sp., Strain Ag1, NR-50119; *Acinetobacter* sp., Strain Ag1, NR-50121; *Serratia* sp., Strain Ag2, NR-50123; *Elizabethkingia anophelis*, Strain Ag1, NR-50124; and *Cedecea* (*Enterobacter*) sp., Strain Ag1, NR-50125.

We declare that we have no conflicts of interest.

## Materials & Methods

### RESOURCE TABLE

| REAGENT or RESOURCE | SOURCE | IDENTIFIER |
| --- | --- | --- |
| **Biological Samples** |  |  |
| Human Blood | Gulf Coast Regional Blood Center | N/A |
| *P. falciparum* 3D7 | BEI Resources Repository | NIAID, NIH, MRA-102 |
| *P. berghei* ANKA infected *Anopheles stephensi* mosquitoes | NYU Langone Medical Center Insectary | N/A |
| *Anopheles gambiae* G3 mosquitoes | BEI Resources Repository | NIAID, NIH, MRA-132K |
| *Enterobacter* sp. Ag1 | BEI Resources Repository | NIAID, NIH, NR-50125 |
| *Serratia* sp. Ag2 | BEI Resources Repository | NIAID, NIH, NR-50123 |
| *Serratia* sp. | (Ganley et al., 2018) | N/A |
| *Pseudomonas* sp. Ag1 | BEI Resources Repository | NIAID, NIH, NR-50126 |
| *Acinetobacter* sp. Ag1 | BEI Resources Repository | NIAID, NIH, NR-50121 |
| *Elizabethkingia anophelis* Ag1 | BEI Resources Repository | NIAID, NIH, NR-50124 |
| **Chemicals, Peptides, and Recombinant Proteins** |  |  |
| Serratiochelin A | This Paper | N/A |
| Aerobactin | This Paper | N/A |
| Pyoverdine Ag1 | This Paper | N/A |
| Bisucaberin | (Rütschlin et al., 2017) | N/A |
| Pyochelin I & II | Toronto Research Chemicals | Cat# P840365 |
| Enterobactin | Millipore Sigma | Cat# E3910 |
| **Critical Commercial Assays** |  |  |
| Bright-Glo Luciferase Assay System | Promega | Cat# G6081 |
| CellTiter-Fluor Cell Viability Assay | Promega | Cat# E2620 |
| **Experimental Models: Cell Lines** |  |  |
| Human: HuH7 cells | Laboratory of Peter Sorger | N/A |
| **Experimental Models: Organisms/Strains** |  |  |
| *P. berghei*-Luc; Strain background: ANKA | NYU Insectary | N/A |
| **Software and Algorithms** |  |  |
| GraphPad Prism 7 | GraphPad Software | graphpad.com |
| antiSMASH 4.0 | (Blin et al., 2017) | https://antismash.secondarymetabolites.org/#!/start |
| XCMS Online | (Colin A. Smith et al., 2006) | https://xcmsonline.scripps.edu/landing_page.php?pgcontent=mainPage |
| MassHunter Qualitative Analysis | Agilent Software | https://www.agilent.com/en/products/software-informatics/mass-spectrometry-software |
| MUSCLE | (Edgar, 2004) | http://www.drive5.com/muscle/ |
| MEGA | (Tamura et al., 2011) | https://www.megasoftware.net/ |

### CONTACT FOR REAGENT AND RESOURCE SHARING

Further information and requests for resources and reagents should be directed to and will be fulfilled by the Lead Contact, Emily Derbyshire.

### EXPERIMENTAL MODEL AND SUBJECT DETAIL

#### Parasite strains

*Plasmodium berghei* ANKA strains (*P. berghei*-Luc) were freshly harvested from dissected salivary glands prior to experiments from infected *Anopheles stephensi* mosquitoes purchased from the New York University Langone Medical Center Insectary. Blood-stage parasites, *P. falciparum* 3D7 strain, were acquired from BEI Resources Repository (NIAID, NIH, MRA-102, contributed by Daniel J. Carucci).

#### Cell lines

HuH7 cells (gift from Dr. Peter Sorger) were cultured and maintained in Dulbecco’s Modified Eagle Medium (DMEM) supplemented with L-glutamine (Gibco), 10 % heat-inactivated fetal bovine serum (HI-FBS) (v/v) (Sigma-Aldrich), and 1 % antibiotic-antimycotic (Thermo Fisher Scientific) in a standard tissue culture incubator at 37ºC and 5 % CO_2_.

### METHOD DETAILS

#### General experimental procedures

An Agilent 1200 Series ChemStation Preparative HPLC system equipped with a diode array and a SUPELCO^®^ SUPELCOSIL™LC-18 column (5 μm, 250 × 21.2 mm) was used for the preparative purification of serratiochelin A, aerobactin, and pyoverdine Ag1. HPLC was monitored at 254 nm for the purification of compounds, unless otherwise stated. For all metabolomic studies, LCMS-ESI was conducted on an Agilent 6224 LC/MS-TOF (high resolution) system equipped with a diode array and an Agilent ZORBAX SB-C18 (5 μm, 2.1 × 50 mm) column. For MS/MS, an Agilent 6460 Triple Quadrupole LC-MS was used.

#### Bacterial strains and media

The bacterial strains used in this study are given in **Data Set S1E**. Bacteria were grown in liquid Luria Bertani (LB) medium or solid bacteriological medium with 25 g L^−1^ (pH 7) and Difco Bacto agar at 15 g L ^−1^ (pH 7) at 30ºC, unless otherwise stated. To prepare metal deficient LB broth, 0.5 g L^−1^ of Chelex® 100 sodium form (Sigma-Aldrich) was added and shaken at 250 rpm at 5ºC for 2 hours, followed by filtering of the resin and subsequent pressure sterilization. For cultures grown in M9 minimal media, 0.4 % glucose was used as a carbon source.

#### Generation of phylogenetic tree of bacteria isolated from mosquito microbiomes

16S sequences of the 33 bacterial species previously isolated from mosquito microbiomes were analyzed. Sequences were aligned using MUSCLE (Edgar, 2004) alignment in MEGA (Tamura et al., 2011) to construct a maximum likelihood tree with 100 bootstrap replicates using Nearest-Neighbor-Interchange (NNI) heuristic and Tamura-Nei method.

#### Computational analysis of bacterial BGCs from mosquito microbiomes

Genome sequences of bacteria originally isolated from bacteria were compiled from NCBI and were subject to analysis by ClusterFinder with a minimum of 5 coding sequences (CDSs) and 5 biosynthesis-related PFAM domains per cluster and a ClusterFinder probability of 60 %. The genomes were additionally subject to antiSMASH 4.0 to detect BGCs in known classes. The accession numbers, nucleotide numbers, and probability score for each cluster is included in **Data Set S1B**.

#### Prediction of biological function of metabolites from corresponding BGC and dereplication of clusters

To generate predictions of biological functions of metabolites corresponding to each antiSMASH cluster, each cluster was compared to previously characterized clusters. As an initial search within the antiSMASH program, investigation of homologous known gene clusters was utilized to identify similar clusters that are already deposited within the Minimum Information about a Biosynthetic Gene Cluster (MIBiG) repository (Medema et al., 2015). Clusters with genes with similarity to over 50 % of MIBiG standard clusters were further analyzed. Specifically, biosynthetic CDSs from the query clusters were analyzed against the MIBiG cluster CDSs via protein BLAST (Altschul et al., 1997) to assign putative CDS functions. Query clusters that did not have clear matches for MIBiG reference clusters were analyzed through BLASTp and subsequent literature searches. Specifically, core biosynthetic CDSs were analyzed by BLASTp to find the closest related characterized enzymes/BGCs that are not deposited within the MIBiG repository. For non-modular BGCs (all except for type I PKS & NRPS BGCs), matching homologs from previously characterized clusters to all core biosynthetic genes of the mosquito clusters (excluding building block genes and accessory genes) was necessary to assign a putative function assignment. Variation in additional non-core biosynthetic genes was allowed. For modular-based BGCs, like type I PKS and NRPS, domain architecture was analyzed. Conservation in overall domain architecture and all core biosynthetic genes was necessary for putative function prediction assignment, with the exception of truncated contigs (See Clusters Serratia70, Serratia99, Serratia130 & Serratia151 in **Data Set S2**). Slight variations in monomer incorporations for acyltransferase (for PKS BGCs) and adenylation (for NRPS BGCs) were tolerated, as well as additional epimerase domains within NRPS BGCs. The structure of the molecules from clusters Chromobacterium4 and Chromobacterium6 were previously investigated (Saraiva et al., 2018), therefore no phylogenetic analysis was conducted for these two gene clusters and predicted functions were based off of the corresponding metabolites. Summary of predicted functions are found in **Data Set S1C**. Detailed bioinformatic analysis to predict functions are found in **Data Set S2**.

In an attempt to understand the chemical diversity of molecules produced by BGCs detected by antiSMASH, highly similar biosynthetic gene clusters were dereplicated. To meet this criterion, all biosynthetic genes need to be conserved across the two clusters with over 85 % protein coverage and over 50 % protein identity of each CDS. For modular BGCs, predictions of monomer incorporations additionally need to be conserved in order to be considered a replicate. Nearly all replicate BGCs were found in bacteria within the same genus. It is important to note that some BGCs exist on separate genetic loci and function together to produce metabolites, as seen in pyoverdine biosynthesis (Ravel and Cornelis, 2003), or may be interrupted at the end of contig sequences; thus, the predicted total number of BGCs may overestimate the actual BGCs. In instances of disparate predicted BGCs that are well-established to work together or could be identified on separate contigs, these BGCs were counted as one in the total dereplicated BGCs (Pseudomonas10 + Pseudomonas24 and Pseudomonas1 + Pseudomonas5). Replicate BGCs are summarized in **Data Set S1D**.

#### Optimization of siderophore production and detection in mosquito microbiome species

In order to conduct differential metabolomics to identify siderophore products, two culturing conditions for each strain were optimized. The first condition had zero to little siderophore production, while the second successfully produced siderophore product(s). To monitor siderophore production, supernatant aliquots were taken at various time points during growth and tested via the liquid chromo azurolsulfonate (CAS) colorimetric assay (Schwyn and Neilands, 1987). The following strains were able to grow well in M9 minimal media: *Serratia* sp., *Enterobacter* sp. Ag1, and *Pseudomonas* sp. Ag1. For each of these strains, growth overnight at 30ºC in M9 minimal media without additional supplementation of iron, resulted in CAS assays color change indicating siderophore production. With the supplementation of FeCl_3_ at 1 mM, siderophore production was halted or significantly decreased as indicated by the CAS assay. For *Serratia* sp. Ag2, growth in metal deficient LB broth for 3–4 days resulted in production of enterobactin. *E. anophelis* Ag1 produced the greatest quantities of bisucaberin when grown in metal deficient media with peptone (1.0 wt %), yeast extract (0.5 wt %), glucose (0.4 wt %), Na_2_HPO_4_ (6.7 g/L), KH_2_PO_4_ (3 g/L), NH_4_Cl (1.67 g/L), NaCl (0.5 g/L), MgSO_4_ (200 mM), Ca_2_Cl (100 mM). *Acinetobacter* sp. Ag1 did not grow in metal deficient media, however acinetoferrin was still produced in normal LB liquid media.

#### Extraction and LCMS-ESI analysis of mosquito microbiome metabolites

All bacterial species were streaked out on LB agar and growth overnight at 30ºC. Colonies were subsequently picked and grown in 5 mL of various optimized iron-deficient or iron-supplemented media for various times at 30ºC at 250 rpm. For each sample, 3 biological replicates were used. After growth, cultures were pelleted, and the supernatants were freeze-dried. The lyophilized supernatants were dissolved in 3 mL of MeOH and vortexed (2 × 0.5 minute), pelleted, and the MeOH layer was dried *in vacuo*. The MeOH extracts were reconstituted in 150 μL of 50:50 H_2_O:MeCN and filtered. The resulting solution was diluted 1:10 in MeCN and used for metabolomic studies.

#### Differential metabolomics of microbiome species with various levels of iron

Samples were analyzed by reversed-phase chromatography on an Agilent 6224 LCMS-TOF using an Agilent ZORBAX SB-C18 (5 μm, 2.1 × 50 mm) column. For each sample, the following mobile phases were used: 98:2 (v:v) H_2_O:MeCN with 0.3 % formic acid (A) and 98:2 (v:v) MeCN:H_2_O with 0.3 % formic acid (B) with a flow rate of 0.350 mL min^−1^, with the following method: 0–5 minutes, isocratic 100 % A; 5–20 minutes, linear gradient from 0–100 % B; 20–30 minutes, isocratic 100 % B; 30–32 minutes, linear gradient from 0–100 % A; from 32–40 minutes, isocratic 100 % A. All MS data was collected in positive ion mode under the following parameters: mass range: 100-1100 m/z; drying gas: 325°C, 11 L/min; nebulizer: 33 psig; capillary: 3500 V; fragmentor: 175 V; skimmer: 65 V; OCT 1 RF Vpp: 750 V; 1000 ms per spectrum. The initial MS data were analyzed by MassHunter Qualitative Analysis software (Agilent). Subsequent differential metabolomic analysis was carried out with XCMS using pairwise analysis comparing the iron-deficient runs against the iron-supplemented runs with three biological replicates of each (Colin A. Smith et al., 2006). Data from each analysis is included in **Data Set S3**, which includes a list of mass peaks (*m*/*z* values), retention times, peak intensities, fold change, log2[fold change], *P* values (two-tailed unequal Student’s *t*-test), and −log[*P*] values. To visualize volcano plots, log2[fold change] was plotted against −log[*P*] in GraphPad Prism.

#### Production and isolation of serratiochelin A, aerobactin, and pyoverdine Ag1

##### Serratiochelin A

Serratiochelin A was isolated as previously described (Seyedsayamdost et al., 2012) with slight modifications. Briefly, *Serratia* sp. was streaked out on LB agar grown overnight and then picked and grown in 5 mL of M9 minimal media and grown at 30ºC at 250 rpm overnight. The overnight culture was used to inoculate four 4 L unbaffled Fernbach flasks each with 1 L of M9 minimal media. The cultures were grown at 30ºC/250 rpm for 4 days. The resulting culture was spun down and the supernatants were extracted with EtOAc (1 L per flask) and washed with brine, dried over Na_2_SO_4_, and concentrated *in vacuo*. The crude extract was reconstituted in 5 mL of MeCN and fractionated by preparative reversed-phased HPLC using the following gradient, monitoring at 254 nm: 0–10 minutes, isocratic 100% H_2_O; 10–15 minutes, linear gradient from 0% MeCN to 30% MeCN; 15–40 minutes, linear gradient from 30% MeCN to 60% MeCN; 40–50 minutes, linear gradient from 60% MeCN to 100% MeCN; 50–55 minutes, isocratic 100% MeCN. Since serratiochelin A is labile to acid hydrolysis, the HPLC solvents were not acidified. Fractions were analyzed by HR-ESI and fractions solely containing the serratiochelin A mass were pooled, concentrated, and lyophilized to afford purified serratiochelin A (5–10 mg).

##### Aerobactin

Aerobactin was purified via a modified protocol from Holt & Butler (Haygood et al., 1993). Briefly, *Enterobacter* sp. Ag1 was streaked out on LB agar grown overnight and then picked and grown in 5 mL of M9 minimal media and grown at 30ºC at 250 rpm overnight. The overnight culture was used to inoculate four 4 L unbaffled Fernbach flasks each with 1 L of M9 minimal media. The cultures were grown at 30ºC at 250 rpm for 4 days. The resulting culture was pelleted, and the supernatants were collected. To the supernatants 20 g L ^−1^ of Dowex ® 1X8 chloride form (100–200 mesh, Sigma-Aldrich) resin was added and shaken at 250 rpm at 5ºC for 24 hours. The resin-slurry was applied to a gravity column and washed with ultrapure H_2_O (3 × 250 mL). The siderophore was eluted with 50 % MeOH:water and fractions containing siderophore activity via the CAS assay were pooled, concentrated, and lyophilized. The resulting freeze-dried pellet was reconstituted in 10 mL of MeOH and fractionated by preparative reversed-phase HPLC using the following gradient, monitoring at 220 nm: 0–10 minutes, isocratic 100% H_2_O; 10–15 minutes, linear gradient from 0% MeCN to 30% MeCN; 15–40 minutes, linear gradient from 30% MeCN to 60% MeCN; 40–50 minutes, linear gradient from 60% MeCN to 100% MeCN; 50–55 minutes, isocratic 100% MeCN. Both HPLC solvents were acidified with 0.01% formic acid. Fractions were analyzed by HR-ESI and fractions solely containing the aerobactin mass were pooled, concentrated, and lyophilized to afford pure aerobactin (6 mg).

##### Pyoverdine Ag1

For isolation of the mixture of pyoverdine Ag1, *Pseudomonas* sp. Ag1 was streaked out on LB agar and grown for 2 days at 30ºC. A colony was picked and grown overnight in LB media and grown overnight at 30ºC at 250 rpm. The overnight culture was used to inoculate four 4 L unbaffled Fernbach flasks each with 1 L of M9 minimal media. The cultures were grown at 28ºC at 200 rpm for 5 days. The resulting culture was centrifuged, and the supernatants were taken. A portion of the supernatants (300 mL) was lyophilized and then extracted with MeOH (300 mL) and dried *in vacuo*. The crude material was fractionated by preparative reversed-phased HPLC using the following gradient, monitored at 405 nm: 0–10 minutes, isocratic 100% H_2_O; 10–15 minutes, linear gradient from 0% MeCN to 30% MeCN; 15–40 minutes, linear gradient from 30% MeCN to 60% MeCN; 40–50 minutes, linear gradient from 60% MeCN to 100% MeCN; 50–55 minutes, isocratic 100% MeCN. Both HPLC solvents were acidified with 0.01% formic acid. Fractions were analyzed by LCMS HR-ESI and fractions solely containing the pyoverdine Ag1 mass were pooled, concentrated, and lyophilized to afford pure pyoverdine Ag1 (0.2 mg).

##### Partial structure elucidation of pyoverdine Ag1

The structure of pyoverdine Ag1 was partially characterized through mass spectrometry and biosynthetic considerations. LCMS-TOF analysis of *Pseudomonas* sp. Ag1 extracts indicated a UV peak absorbing at 405 nm with a *m*/*z* equal to 647.78512 ([M+2H]^2+^). From this, a molecular formula of C_53_H_79_N_15_O_23_ was predicted, which is similar to other characterized pyoverdines. Based on the MS/MS profile of the pyoverdine, along with biosynthetic considerations from the BGC (**Data Set S2**), the peptide sequence along with the chromophore structure were predicted (**Fig. S4**). The peptide sequence (Chromophore-Ser-fhOrn-Lys-Thr-Asn-Gly-Ser-fhOrn) varied slightly from the adenylation predictions from antiSMASH (Chromophore-Ser-fhOrn-Gly-Thr-Ala-Gly-Ser-fhOrn). Absolute configuration of amino acids was predicted based on presence and location of epimerase domains within the pyoverdine BGC. Additionally, our analysis indicates a linear peptide sequence, while similar pyoverdines are cyclized through the C-terminal amino acid and the threonine residue. The linear peptide sequence may be due to hydrolysis during preparation or analysis. Further NMR work would be needed for unambiguous structure assignment.

#### Anti*-Plasmodium* liver-stage assays

Luciferase-expressing *P. berghei* ANKA (NYU Langone Medical Center Insectary Core Facility) harvested from *A. stephensi* salivary glands were used to infect HuH7 hepatocytes as previously described (Derbyshire et al., 2012). Cells were seeded 24 hours before infection at 4,000 cells/well in a 384-well plate. Compounds were added in triplicate in a dose dependent manner 30 minutes prior to infection. As a negative control, 0.1% DMSO was used. HuH7 cell viability and parasite load were determined 44 hours post infection using CellTiter-Fluor (Promega) and Bright-Glo (Promega) reagents, respectively, according to manufacturer’s protocols. An EnVision plate reader (Perkin Elmer), was used to measure relative fluorescence and luminescence. EC_50_ values were calculated by GraphPad Prism from two biological replicates.

#### Anti-*Plasmodium* blood-stage assays

*P. falciparum* 3D7 parasite (MRA-102) was obtained from BEI Resources and cultured as previously described (Radfar et al., 2009; Raphemot et al., 2019). Parasites were synchronized with 5 % D-sorbitol (Sigma) at 37ºC for 10 minutes and adjusted to 2 % parasitemia and 2 % hematocrit prior to the assay. One-hundred μL of the culture was dispensed into each well of a 96-well black plate (Corning) containing 100 μL medium in the presence or absence of siderophores (0–100 μM final concentration) in triplicate. Each well contained 0.5% DMSO. The plate was incubated at 37°C for 72 hours. Cells were lysed with 40 μL lysis buffer (20 mM Tris-HCl, pH 7.5, 5 mM EDTA dipotassium salt dihydrate, 0.16 % saponin, 1.6 % Triton X-100) containing fresh 1x SYBR Green I (ThermoFisher Scientific) at room temperature in the dark for 24 hours (Kato et al., 2016). The relative parasite loads were determined using an EnVision 2105 plate reader (PerkinElmer) at 535 nm with excitation at 485 nm. EC_50_ values were calculated by GraphPad Prism from two biological replicates.

#### Mosquito rearing and assessment of microbiome siderophores on *A. gambiae* fecundity and survival

*A. gambiae* G3 strains were reared under standard conditions as previously described (28ºC +/− 1ºC, 12-hour dark/light cycle) (Kukutla et al., 2013). All mosquitoes were provided 10% sucrose solution with 1% of the siderophore solubilized in DMSO or solely 1% DMSO as a vehicle control. The sucrose solution was changed daily. Mice were provided as a blood source for egg production. Live and dead mosquitoes were monitored and counted every day. After exposure to a blood meal, engorged female mosquitoes were counted and separated from mosquitoes that did not feed. Ovaries of engorged females were dissected three days post blood meal and the number of eggs per mosquito were counted for fecundity assessment. All siderophore-feeding experiments were completed with 100 female *A. gambiae* mosquitoes per siderophore and completed in biological triplicate.

### QUANTIFICATION AND STATISTICAL ANALYSIS

#### Serratiochelin A calibration curve generation and concentration quantification within *A. stephensi* & *A. gambiae*

To generate a calibration curve for serratiochelin A, samples at 1 μg mL^−1^, 100 ng mL^−1^, and 10 ng mL^−1^ were prepared in MeCN and subject to LCMS-ESI using the same method as above. An extracted ion current (EIC) of serratiochelin A (scan width of ± 5 ppm) was generated using Agilent MassHunter Workstation software. The areas under the curves were plotted to generate the linear equation. Three mosquito samples of *A. stephensi* (23 days PBM from NYU insectary), where n = 50 for each, were washed, dried, and ground as previously reported. The grounds were separately extracted with 10 mL of MeOH and concentrated *in vacuo*. The same extraction protocol was followed for samples of *A. gambiae*. The following *A. gambaie* samples were tested: 1) 50 male, 0 day sugar-fed, 2) 50 female, 0 day sugar-fed, 3) 50 male, 3 day sugar-fed, 4) 50 female, 3 day sugar-fed, 5) 50 male, 7 day sugar-fed, 6) 50 female, 7 day sugar-fed, 7) 50 male, 10 day sugar-fed, 8) 50 female, 10 day sugar-fed, 9) 50 female, 1 day PBM, 10) 50 female, 2 day PBM, 11) 50 female, 3 day PBM, 12) 50 female, 4 day PBM. The resulting extract was solubilized in MeCN (1.0 mL for *A. stephensi*, and 0.03 mL for *A. gambiae*) and analyzed under the same HR LCMS-ESI conditions used for all metabolomic studies. At this dilution factor the abundance of the serratiochelin A EIC for *A. stephensi* was within our calibration curve (See **Figure S7** for more details).

#### Statistical Information

GraphPad Prism 7 software was used for data analysis. Results are represented as means +/− the standard error of the means (SEM). Statistical tests were performed using either one-way or two-way analysis of variance (ANOVA) with Dunnett’s multiple comparison tests, two-tailed Student’s t-test, unpaired Student’s t-test, or Mantel-Cox test, where appropriate.

### DATA AND SOFTWARE AVAILABILITY

Data sets of the BGCs and the BGC analysis are available as supplementary materials (See **Data Sets 1–2**). Metabolomic data is available as a supplementary data set (**Data Set 3**). Software used herein are available for either download or online usage. This study did not generate any unique code.

## Supplemental Figures

**Figure S1.**
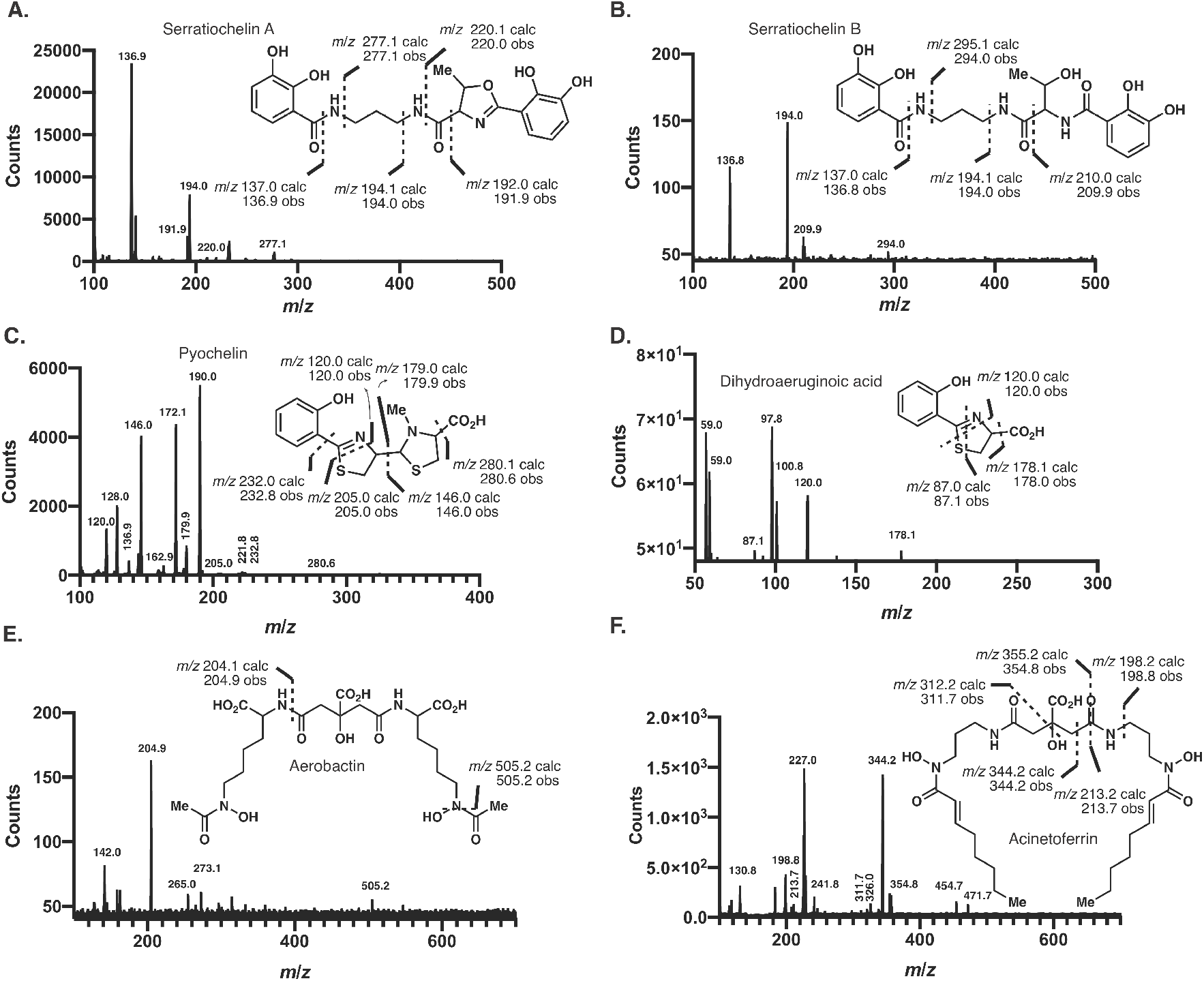
Product Ion Spectra for Mosquito-Microbiome Siderophores. Product ion spectra with predicted and observed fragment ions for (**A**) serratiochelin A, serratiochelin B, (**C**) pyochelin, (**D**) dihydroaeruginoic acid, (**E**) aerobactin, and (**F**) acinetoferrin.

**Figure S2.**
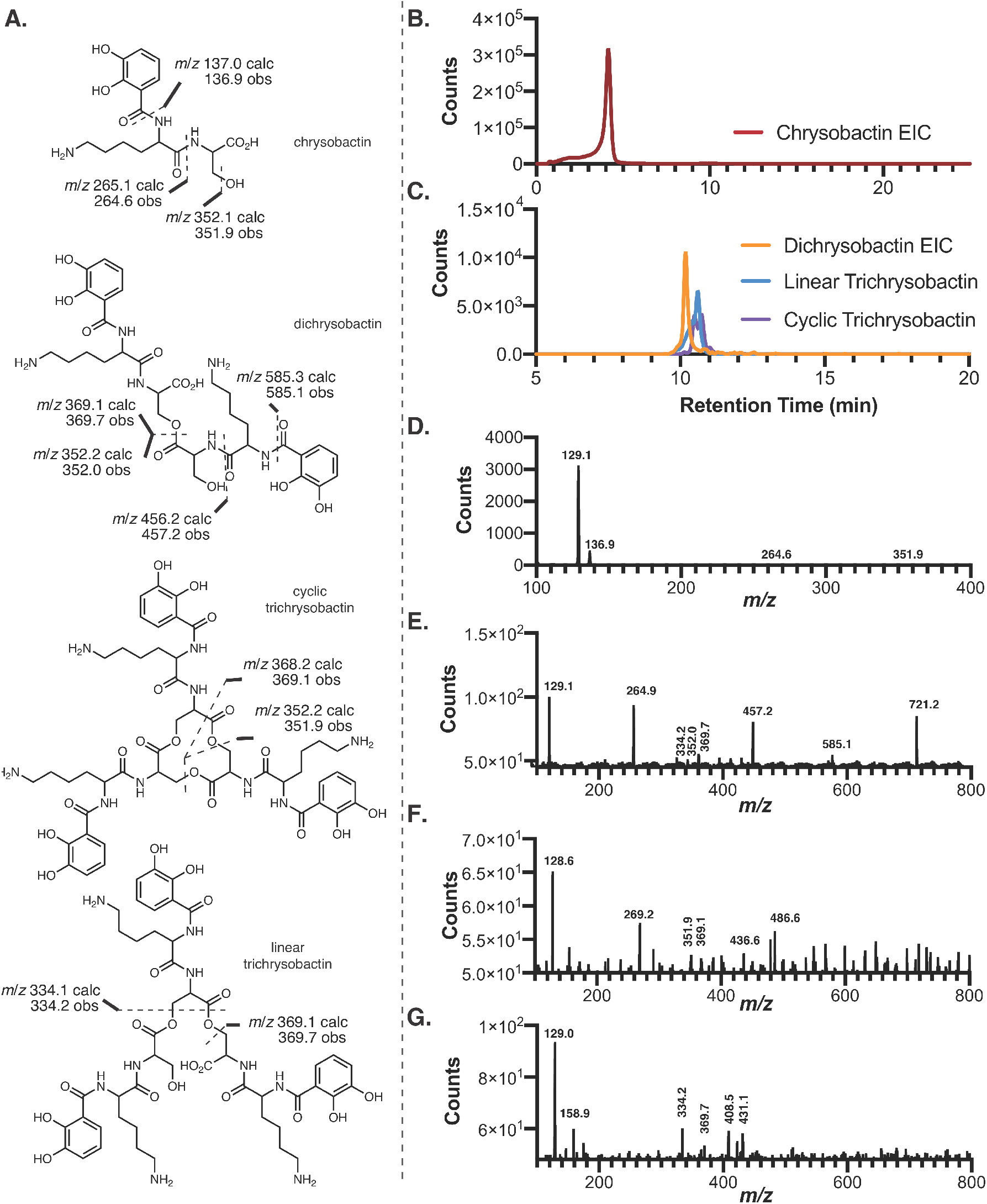
Identification and Confirmation of Production of Chrysobactins by *Serratia* sp. (**A**) Structures of chrysobactin, dichrysobactin, cyclic trichrysobactin, and linear trichrysobactin as well as the predicted and observed fragment ions. (**B**) EIC of chrysobactin from *Serratia* sp. extracts ([M+H] = 370.1609 *m*/*z*). (**C**) EICs of dichrysobactin (orange, [M+H] = 721.3039 *m*/*z*), linear trichrysobactin (blue, [M+H] = 1072.4469 *m*/*z*), and cyclic trichrysobactin (purple, [M+H] = 1054.4364 *m*/*z*). Product ion spectra for (**D**) chrysobactin, (**E**) dichrysobactin, (**F**) cyclic trichrysobactin, and (**G**) linear trichrysobactin.

**Figure S3.**
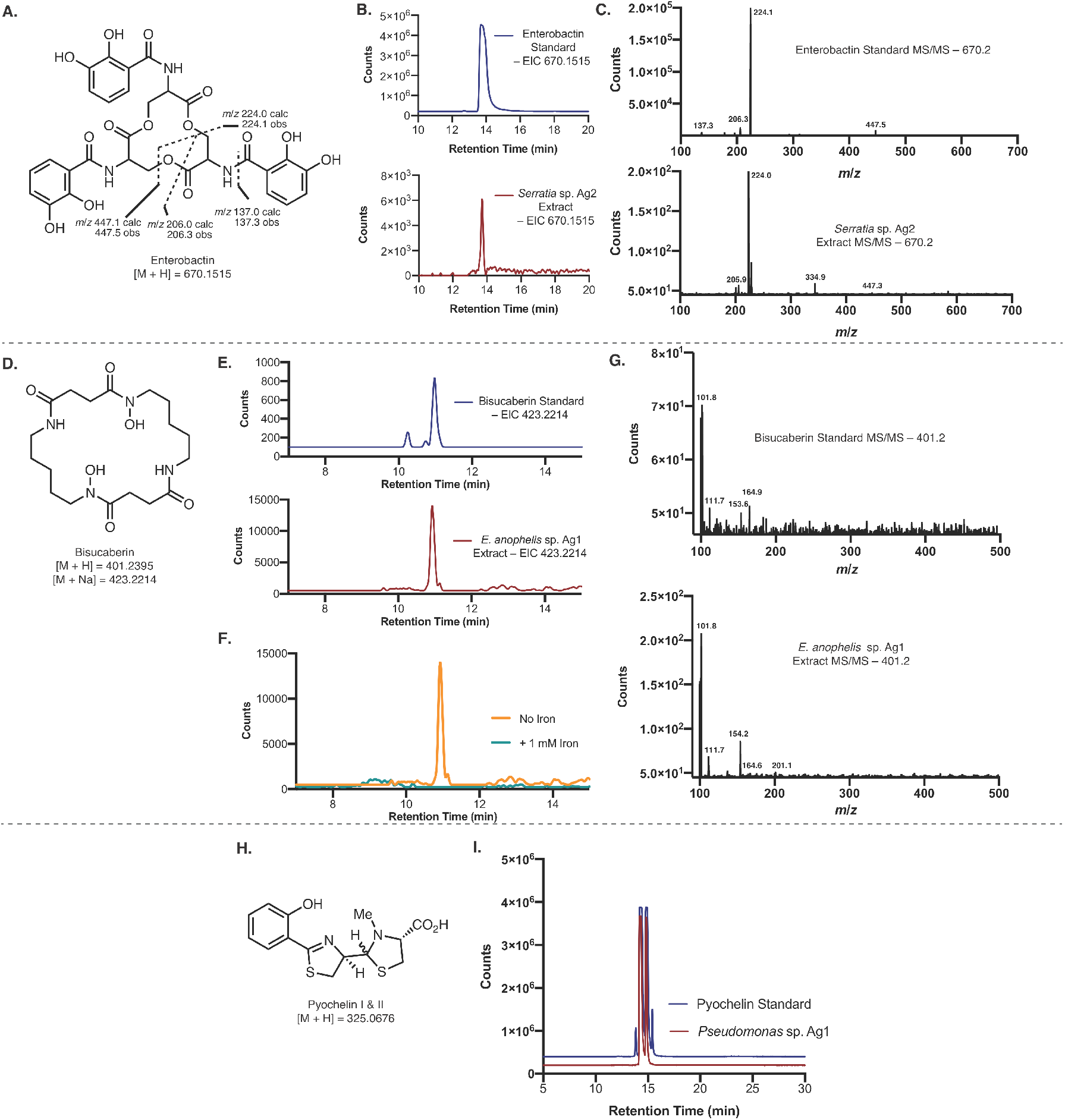
Identification and Confirmation of Production of Enterobactin by *Serratia* sp. Ag2, Bisucaberin by *E. anophelis* Ag1, and Pyochelin by *Pseudomonas* sp. Ag1. (**A**) Structure of enterobactin and the predicted and observed fragment ions. (**B**) EICs of the enterobactin mass ([M+H] = 670.1515 *m*/*z*) from the enterobactin authentic standard (blue) and from *Serratia* sp. Ag2 extracts (red). (**C**) Product ion scan of the enterobactin mass of the enterobactin authentic standard (top) and the *Serratia* sp. Ag2 extracts (bottom). (**D**) Structure of bisucaberin. (**E**) EICs of the bisucaberin mass ([M+Na] = 423.2214 *m*/*z*) from the bisucaberin authentic standard (blue) and from *E. anophelis* Ag1 extracts (red). (**F**) EICs of the bisucaberin from *E. anophelis* extracts when grown without iron (orange) and with 1 mM FeCl_3_ (teal). (**G**) Product ion scan of the bisucaberin mass of the bisucaberin authentic standard (top) and the *E. anophelis* Ag1 extracts (bottom). (**H**) Structure of pyochelin I & II. (**I**) EICs of the pyochelin mass ([M+H] = 325.0676 *m*/*z*) from the pyochelin authentic standard (blue) and from *Pseudomonas* sp. Ag1 extracts (red).

**Figure S4.**
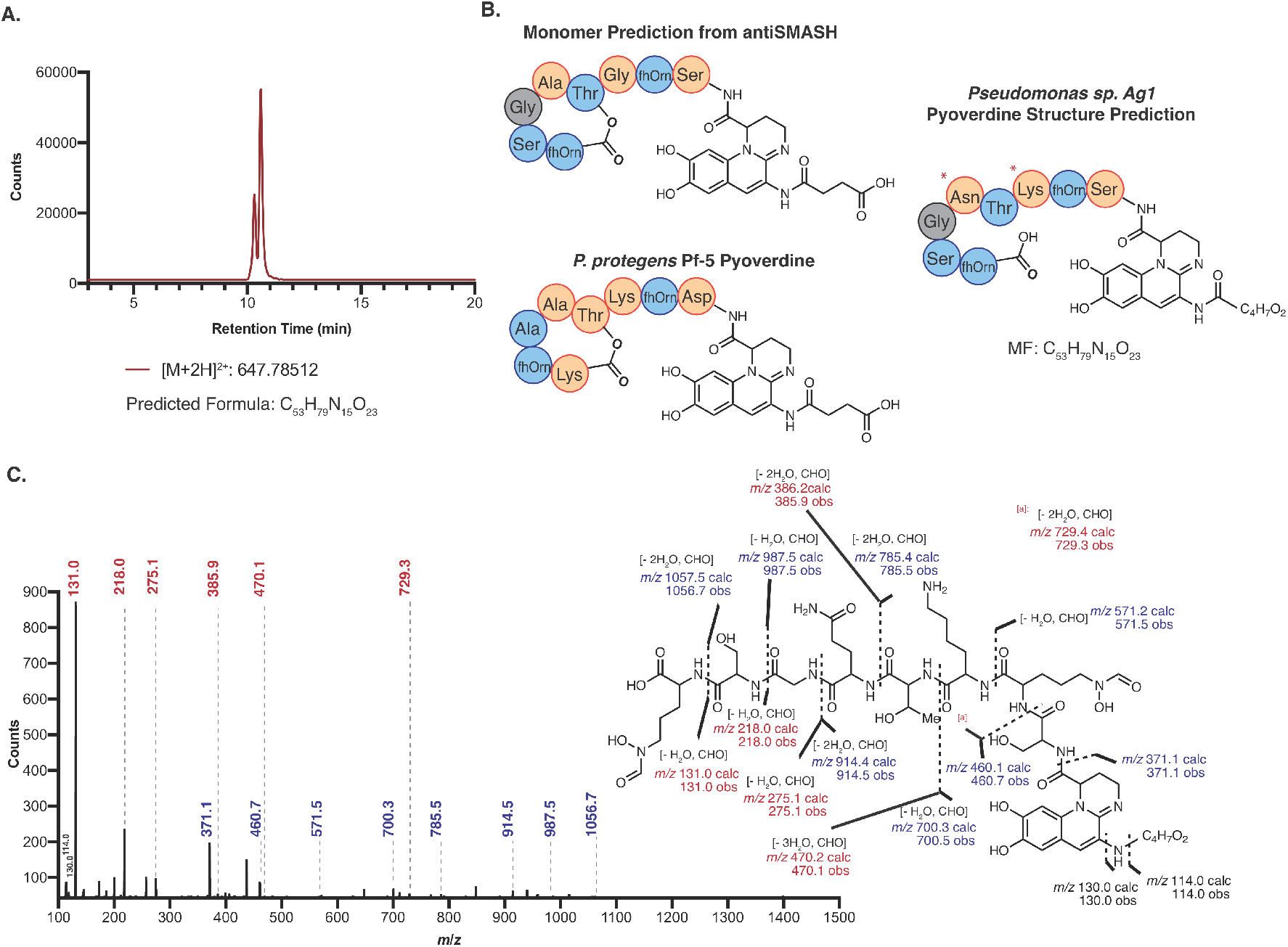
Partial Characterization of Pyoverdine Ag1. (**A**) EIC of predicted [M+2H]^2+^ from *Pseudomonas* sp. Ag1 extracts with the predicted formula of M = C_53_H_79_N_15_O_23_. (**B**) Predicted monomer incorporation of pyoverdine from *Pseudomonas* sp. Ag1 based on antiSMASH predictions, the structure of pyoverdine from the phylogenetically similar *P. protegens* Pf-5 pyoverdine cluster, and the predicted structure based on the tandem mass spectrometry spectra and predicted molecular formula. Orange amino acids represent D-amino acids and blue represent L-amino acids. Amino acid stereochemistry for the pyoverdine Ag1 structures is based on predictions from epimerase domains within the pyoverdine BGC. (**C**) Product ion spectrum of the doubly charged pyoverdine species with the observed and predicted fragment ions shown. Ions in blue represent b-ions and ions in red represent y-ions.

**Figure S5.**
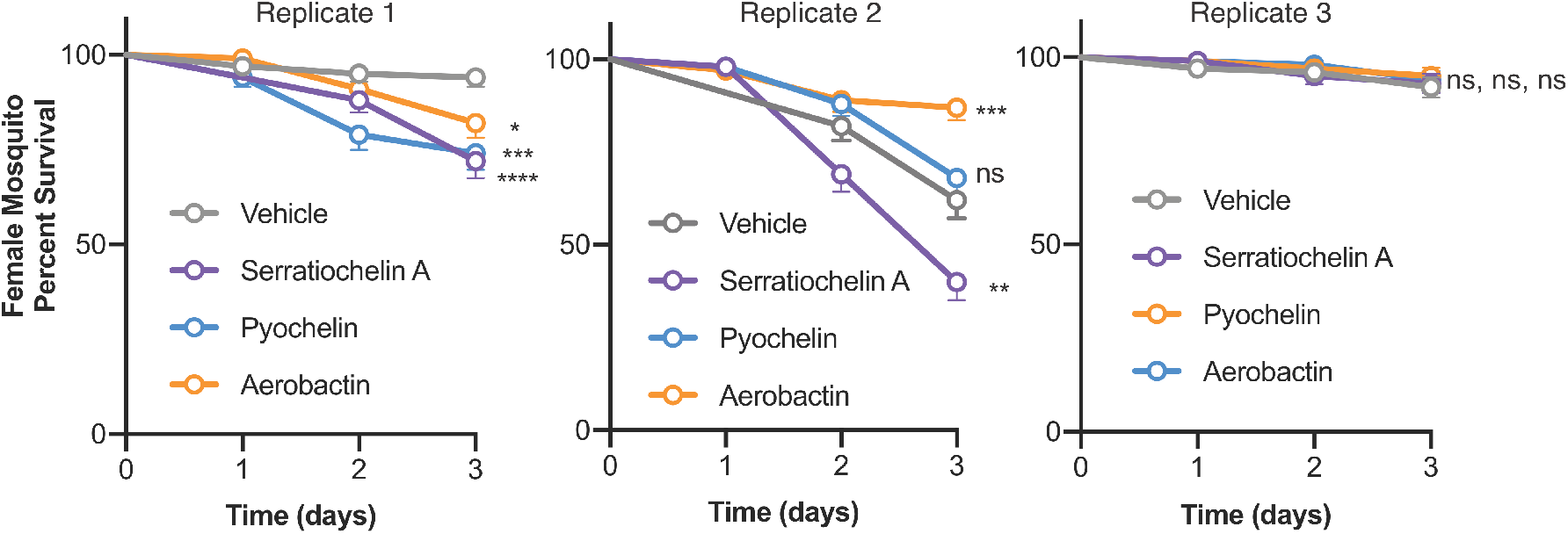
Mosquito Survival Curves with Siderophore Supplementation. (**A**) Survival curves of female mosquitoes (t = 1–3 days) supplemented with siderophores (100 μM) before a blood meal. The three panels represent the data from three separate biological replicates (n = 100 mosquitoes each). The error bars represent the mean +/− the standard error (Log-rank (Mantel-Cox) test, ns = not significant, * *P* < 0.05, ** *P* < 0.01, *** *P* < 0.001 **** *P* < 0.0001).

**Figure S6.**
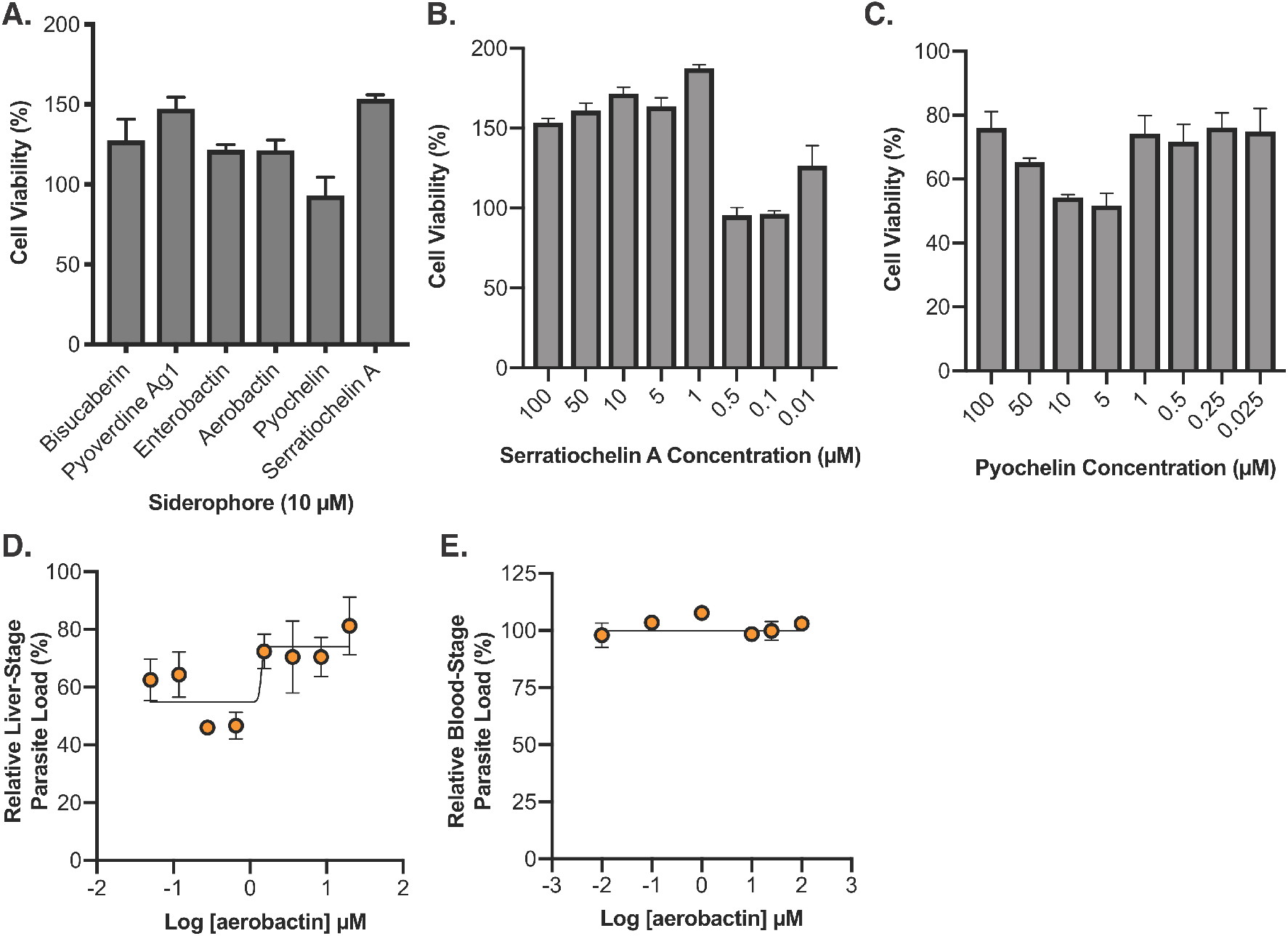
Cell Viability Evaluation of Siderophores and Antimalarial Dose-Down Curves for Aerobactin. (**A**) Cell viability of microbiome siderophores at 10 μM against HuH7 cells normalized to the vehicle control. Cell viability dose-down of (**B**) serratiochelin A and (**C**) pyochelin against HuH7 cells. Dose-down analysis of aerobactin against (**D**) *P. berghei* ANKA parasite load in HuH7 cells and (**E**) *P. falciparum* 3D7 parasite load in human RBCs.

**Figure S7.**
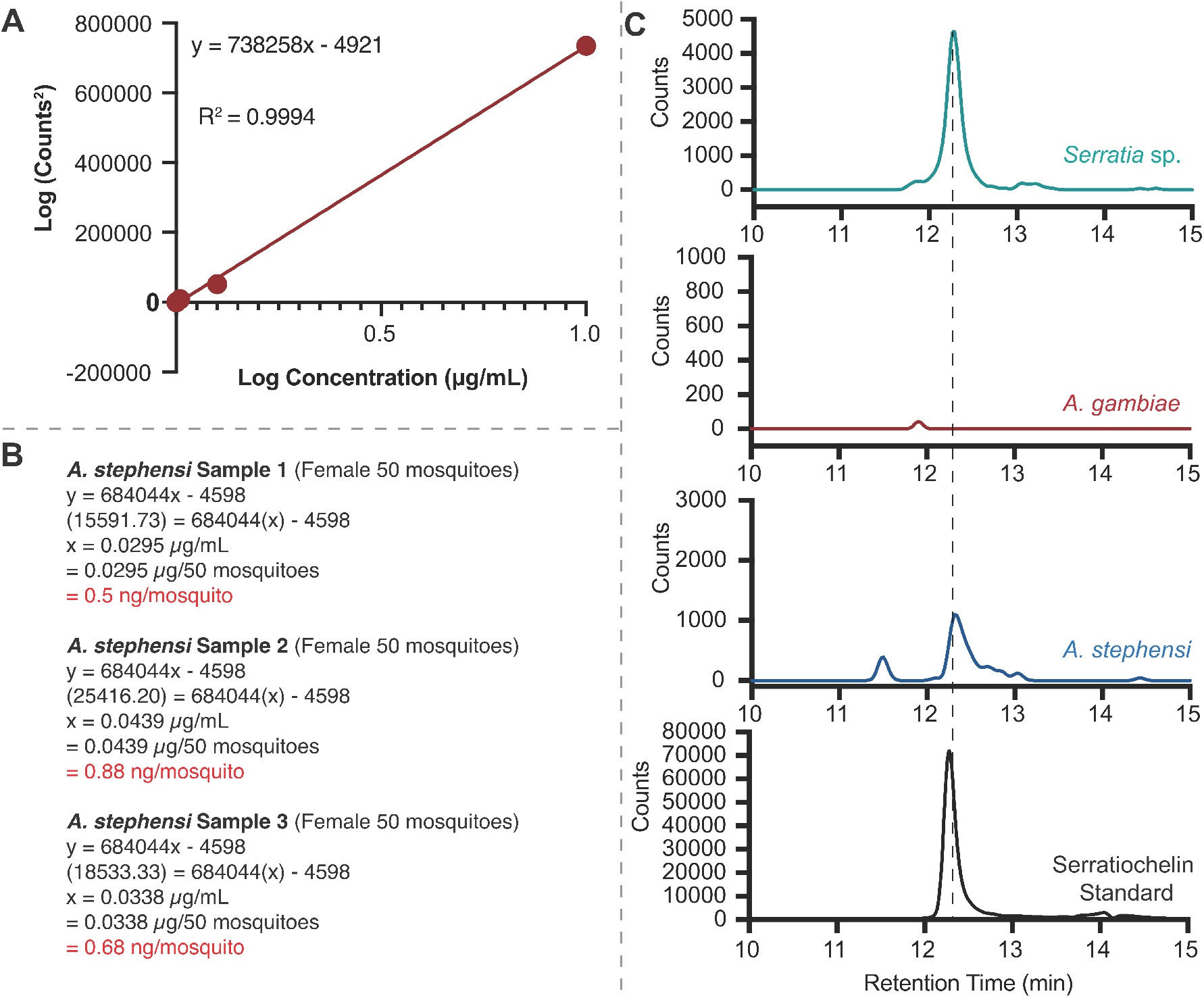
Determination of Serratiochelin A Concentration in Mosquitoes. (**A**) Calibration curve for serratiochelin A. (**B**) Calculations to determine serratiochelin A amount per mosquito in three separate lab-reared *A. stephensi* samples. (**C**) EIC of the serratiochelin A *m*/*z* from *Serratia* sp. culture extracts, *A. gambiae* female mosquitoes, *A. stephensi* female mosquitoes, and a serratiochelin A standard.

## Additional Files

**Supplementary Data Set 1.** (**A**) Bacterial strains, isolation source, and number of BGCs from bioinformatic analysis. (**B**) Accession numbers, nucleotide locations, BGC classifications, and CF probability of all BGCs. (**C**) Functional prediction of aS BGCs. (**D**) Dereplicated/duplicated aS BGCs. (**E**) Strains used in this study.

**Supplementary Data Set 2.** Detailed bioinformatic analysis to determine closest characterized BGC of aS clusters with predicted functions.

**Supplementary Data Set 3.** Data for differential metabolomics for *Serratia* sp., *Pseudomonas* sp. Ag1, and *Enterobacter* sp. Ag1 grown in depleted and high iron.

