## Supplementary Data Set 2 for "A Systematic Analysis of Mosquito-Microbiome Biosynthetic Gene Clusters Reveals Antimalarial Siderophores that Reduce Mosquito Reproduction Capacity"

#### Key:

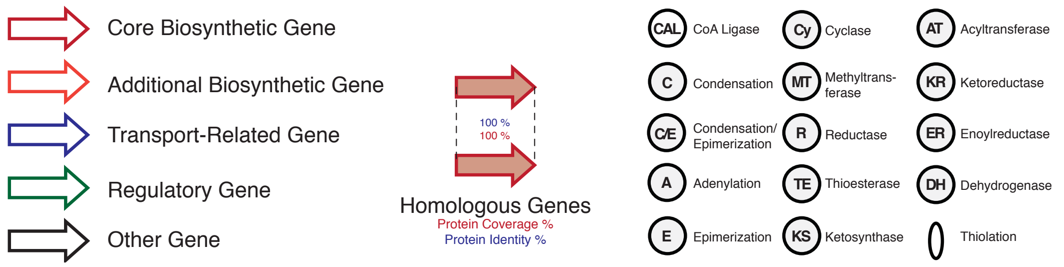


#### Siderophores:

##### Acinetobacter18 & Acinetobacter37:

**Reference**: Funahashi T, Tanabe T, Maki J, Miyamoto K, Tsujibo H, Yamamoto S. 2013. Microbiology 159:678–690.

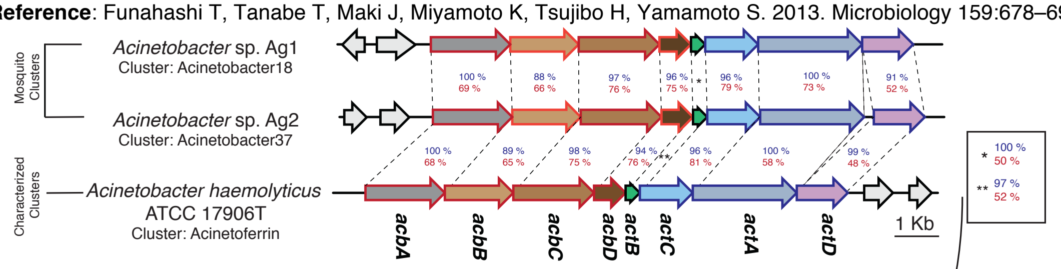


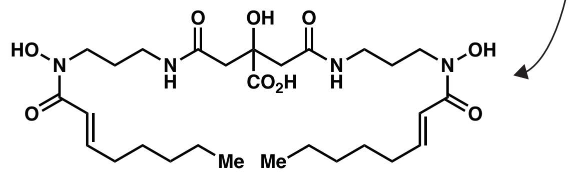


**Acinetoferrin**

##### Elizabethkingia6, Elizabethkingia25, Elizabethkingia46 & Elizabethkingia74:

**Reference**: Thode SK, Rojek E, Kozlowski M, Ahmad R, Haugen P. 2018. PLoS One 13:e0191860.

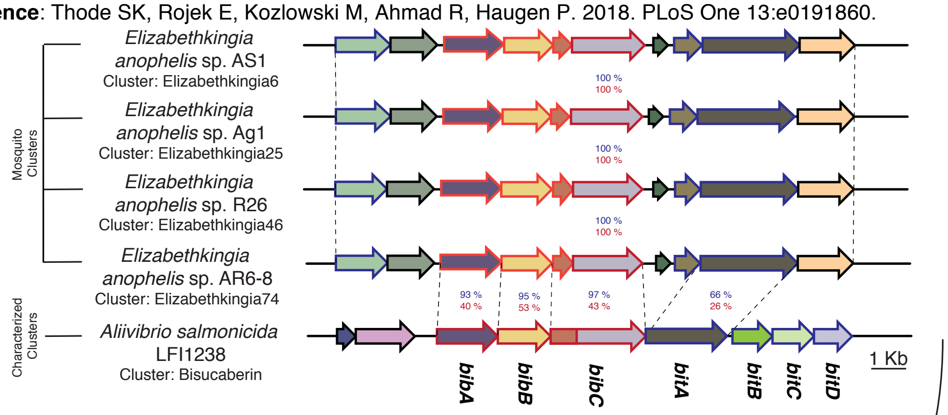


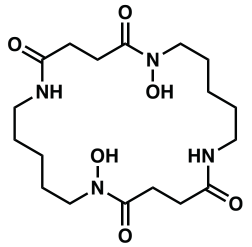


**Bisucaberin**

#### Chromobacterium26:

**Reference**: Mercado-Blanco J, van der Drift KM, Olsson PE, Thomas-Oates JE, van Loon LC, Bakker PA. 2001. J Bacteriol 183:1909–20.

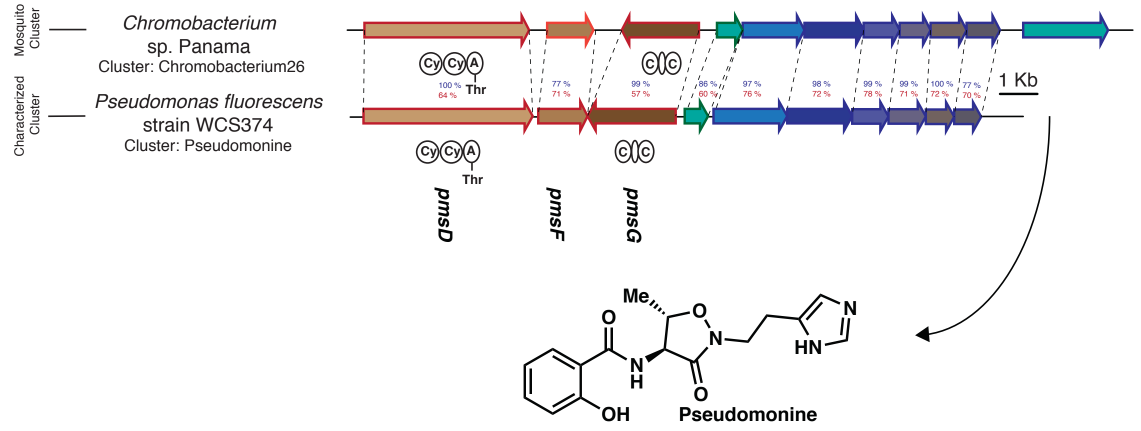


#### Kosakonia4:

**Reference**: Rusnak F, Sakaitani M, Drueckhammer D, Reichert J, Walsh CT. 1991. Biochemistry 30:2916–2927.

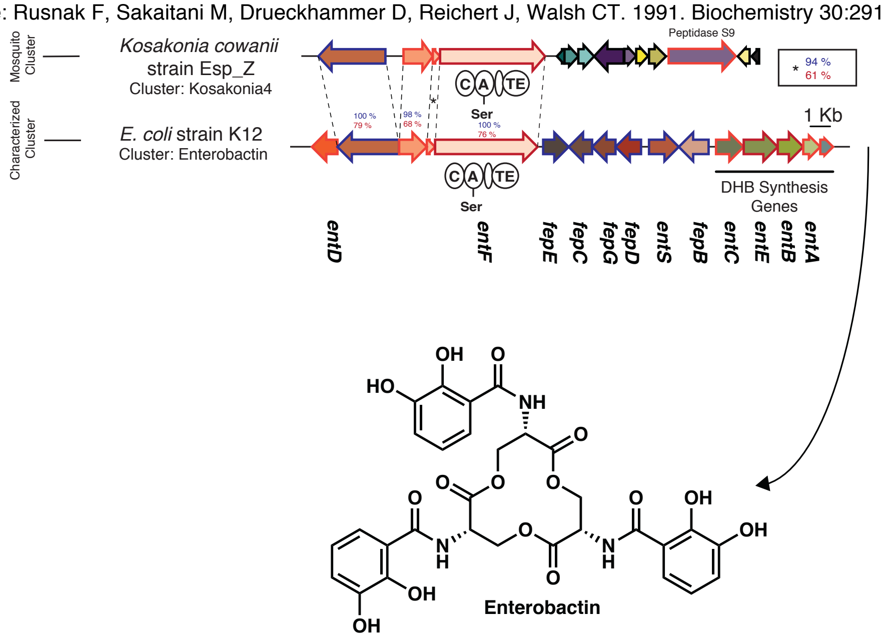


#### Enterobacter1 & Kosakonia13:

**Reference**: Suzuki K, Tanabe T, Moon Y-H, Funahashi T, Nakao H, Narimatsu S, Yamamoto S. 2006. Res Microbiol 157:730–740.

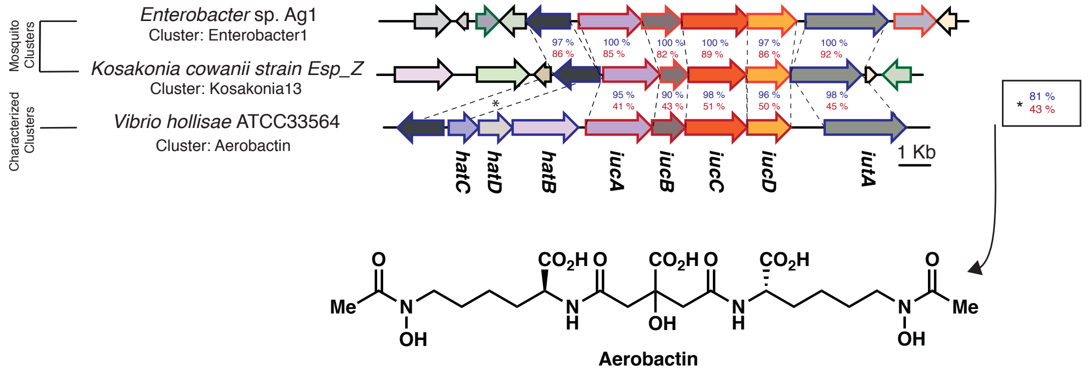

#### Pantoea7:

**Reference**: Smits THM, Duffy B. 2011. Arch Microbiol 193:693–699

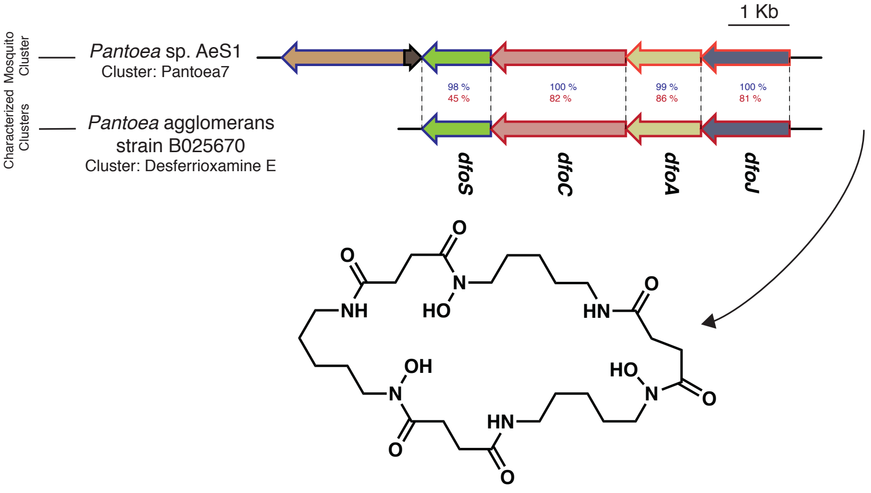


**Nocardamine/Desferrioxamine E**

#### Pseudomonas5 & Pseudomonas24:

**Reference**: Hartney SL, Mazurier S, Girard MK, Mehnaz S, Davis EW, Gross H, Lemanceau P, Loper JE. 2013. J Bacteriol 195:765–76.

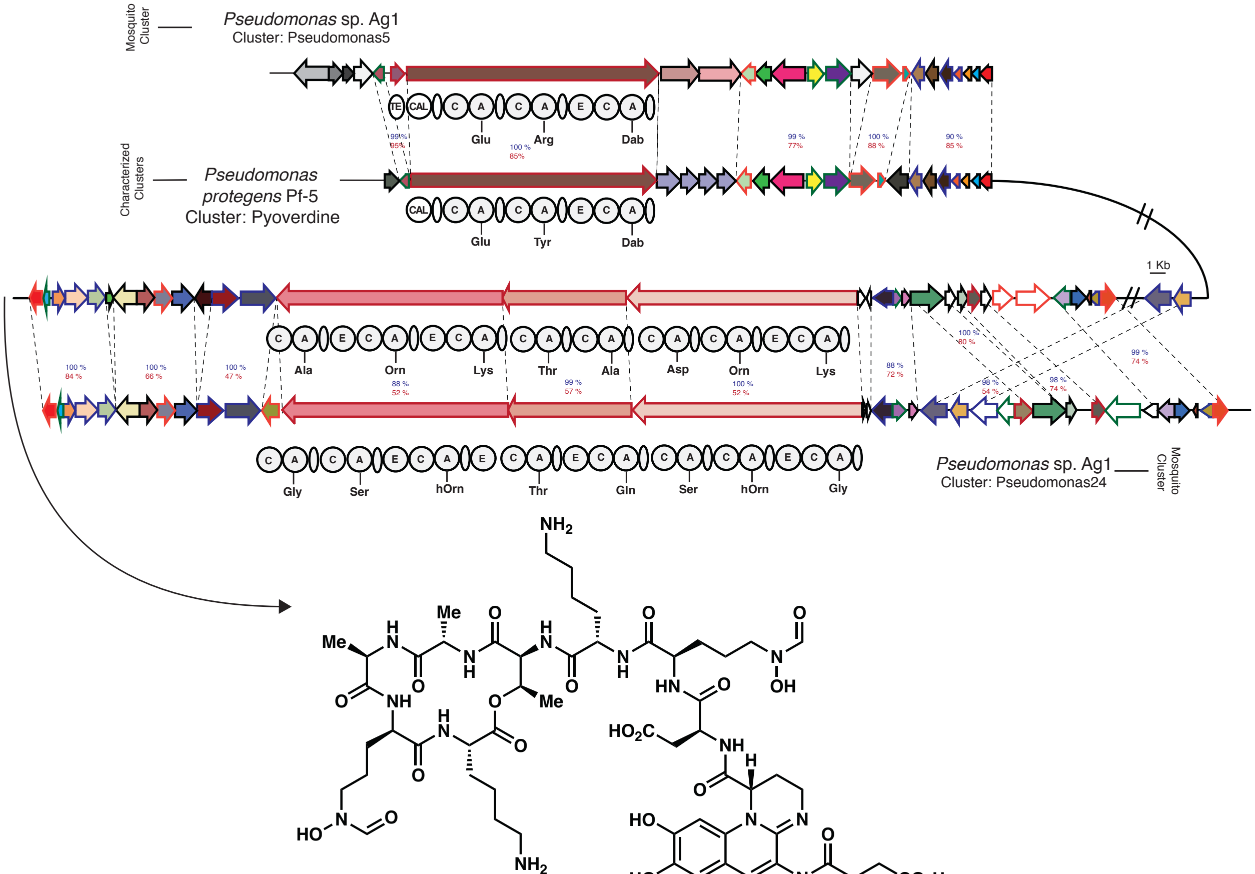


**Pyoverdine Pf5**

#### Pseudomonas36:

**Reference**: Youard ZA, Mislin GLA, Majcherczyk PA, Schalk IJ, Reimmann C. 2007. J Biol Chem 282:35546–35553.

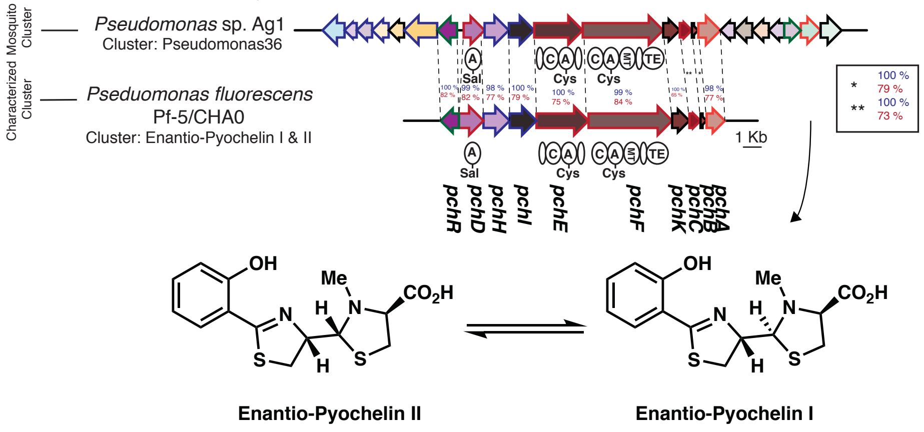


#### Pseudomonas57:

**Reference**: Tunca S, Barreiro C, Sola-Landa A, Coque JJR, Martín JF. 2007. FEBS J 274:1110–1122.

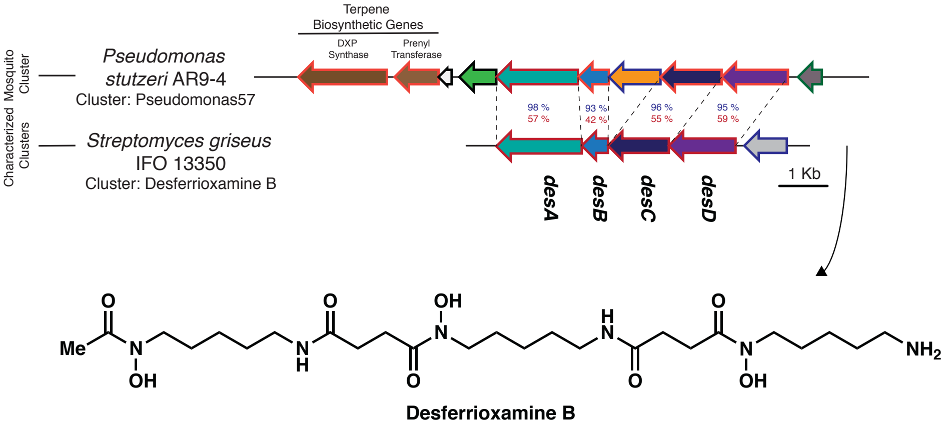


#### Serratia25:

**References**: (a) Seyedsayamdost MR, Cleto S, Carr G, Vlamakis H, João Vieira M, Kolter R, Clardy J. 2012. J Am Chem Soc 134:13550–13553. (b) Suzuki K, Tanabe T, Moon Y-H, Funahashi T, Nakao H, Narimatsu S, Yamamoto S. 2006. Res Microbiol 157:730–740.

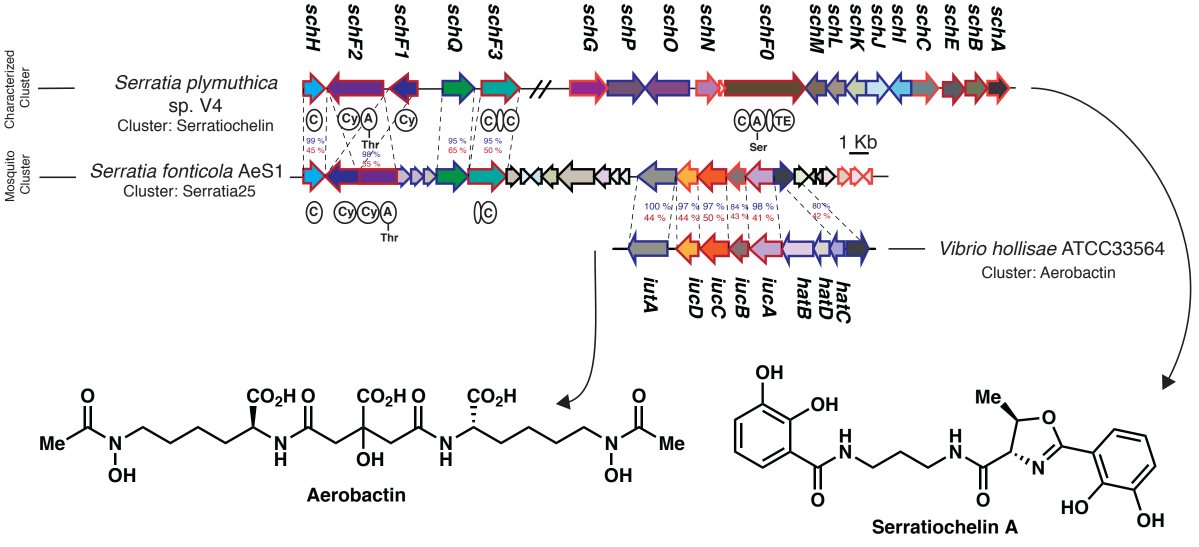

#### Serratia64, Serratia106, & Serratia137:

**Reference**: Seyedsayamdost MR, Cleto S, Carr G, Vlamakis H, João Vieira M, Kolter R, Clardy J. 2012. J Am Chem Soc 134:13550–13553.

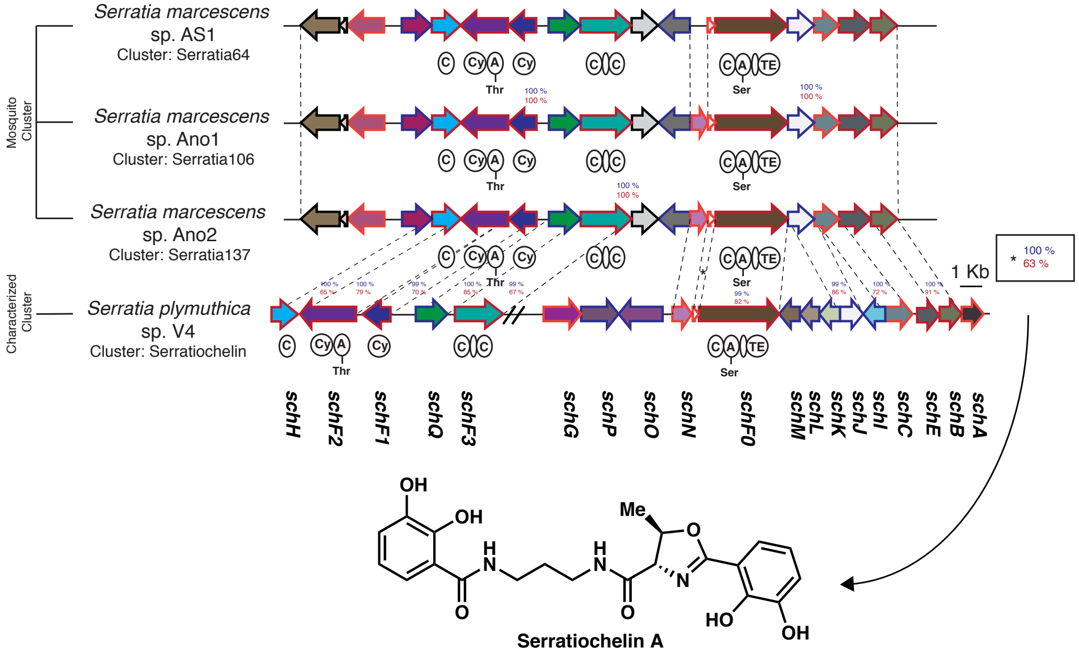


#### Serratia62, Serratia178 & Serratia33:

**Reference**: Rusnak F, Sakaitani M, Drueckhammer D, Reichert J, Walsh CT. 1991. Biochemistry 30:2916–2927.

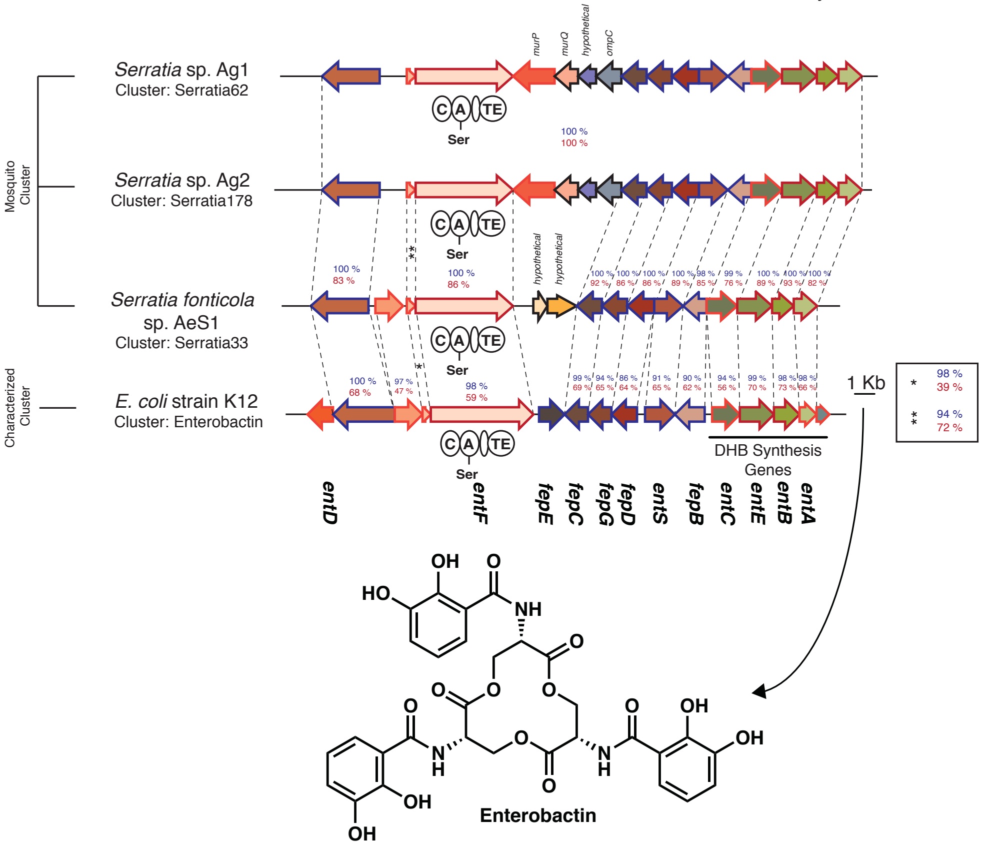

Serratia72, Serratia95 & Serratia126:

**Reference**: Suzuki K, Tanabe T, Moon Y-H, Funahashi T, Nakao H, Narimatsu S, Yamamoto S. 2006. Res Microbiol 157:730–740.

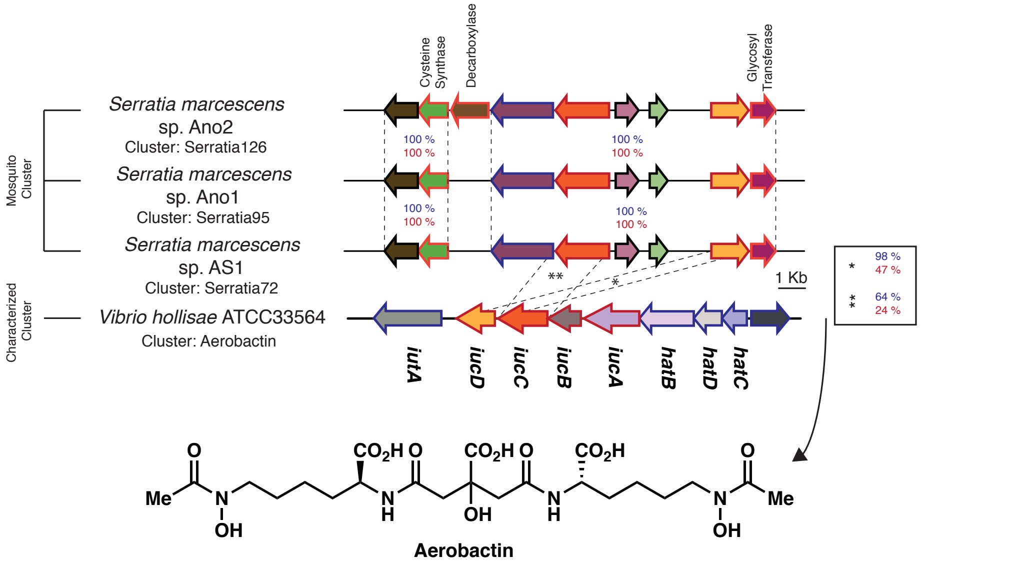


Serratia116, Serratia147 & Serratia81:

**Reference**: Reitz ZL, Sandy M, Butler A. 2017. Metallomics 9:824–839.

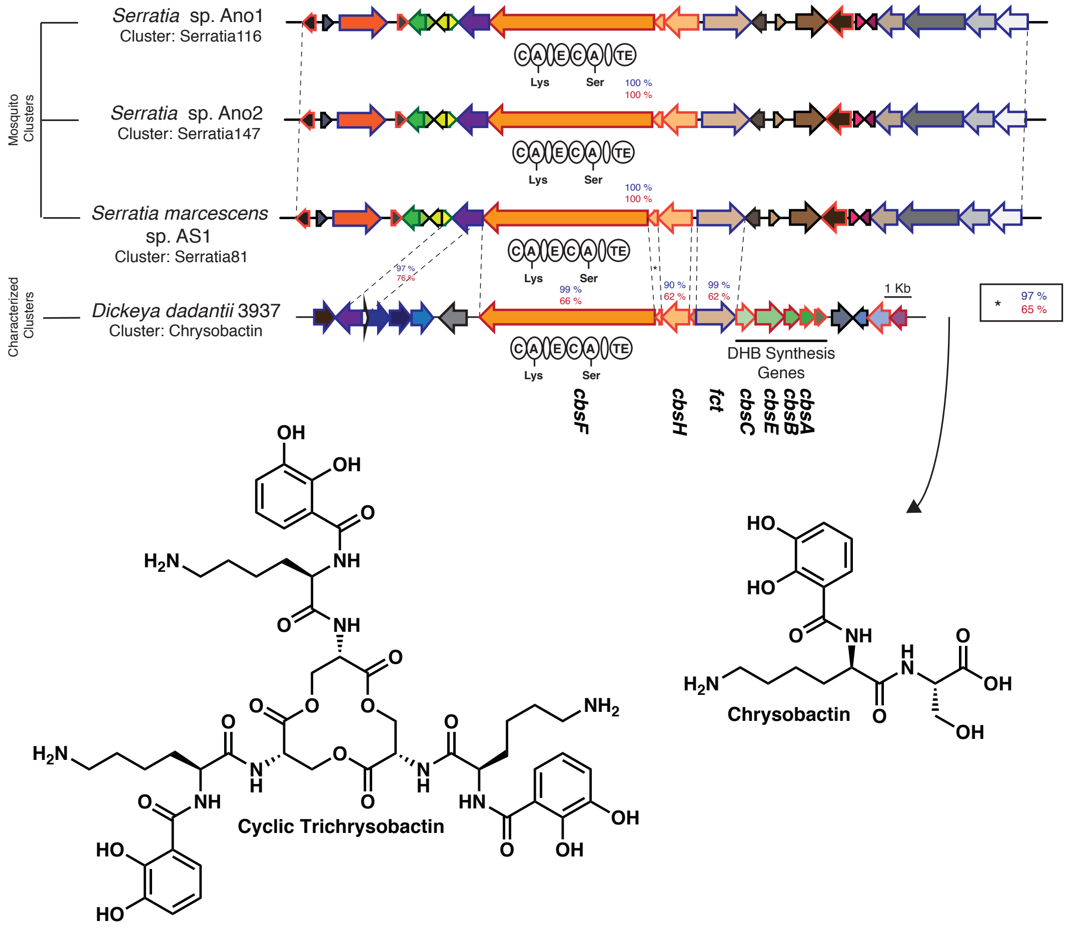

#### Staphylococcus3:

**Reference**: Cheung J, Beasley FC, Liu S, Lajoie GA, Heinrichs DE. 2009. Mol Microbiol 74:594–608.

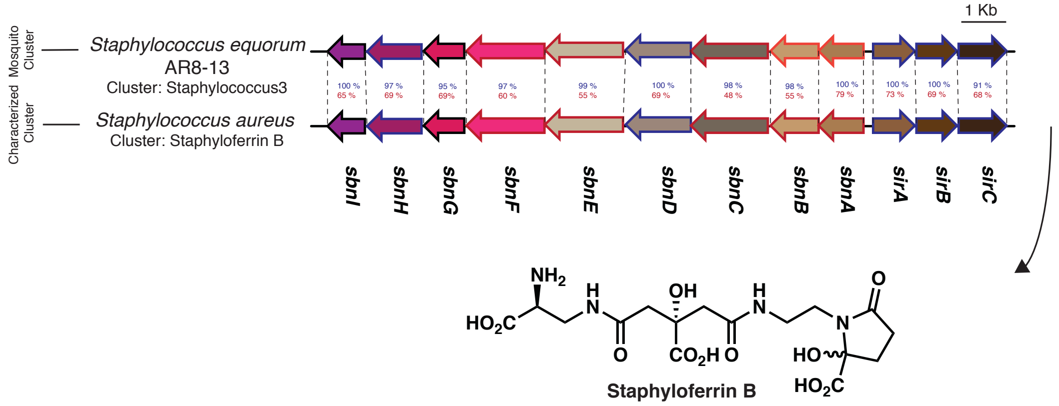


#### Staphylococcus16, Staphylococcus20, Staphylococcus28 & Staphylococcus36:

**Reference**: Cotton JL, Tao J, Balibar CJ. 2009. Biochemistry 48:1025–1035.

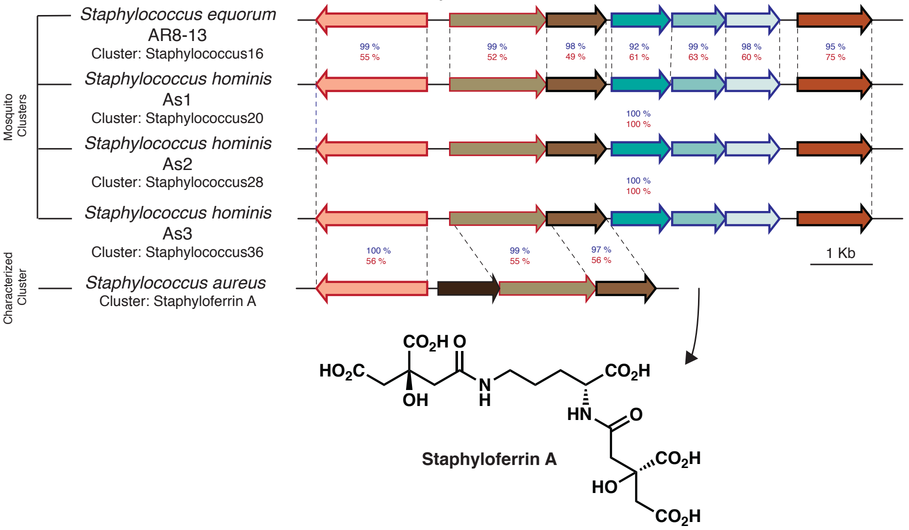


#### Stenotrophomonas9:

**Reference**: Rusnak F, Sakaitani M, Drueckhammer D, Reichert J, Walsh CT. 1991. Biochemistry 30:2916–2927.

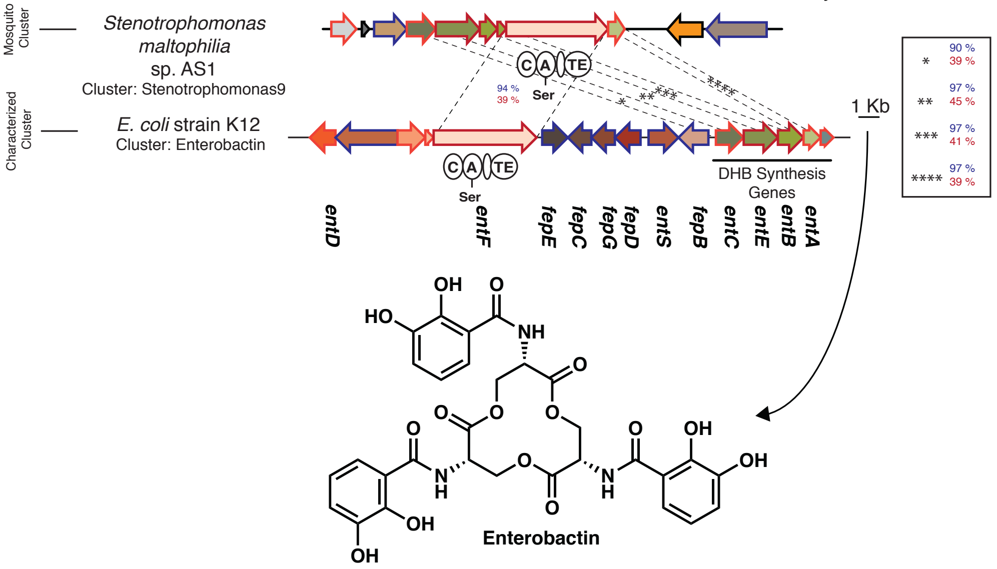

**Pigments:**

Elizabethkingia16, Elizabethkingia26, Elizabethkingia50, & Elizabethkingia75:

**Reference**: Tao L, Yao H, Kasai H, Misawa N, Cheng Q. 2006. Mol Genet Genomics 276:79–86.

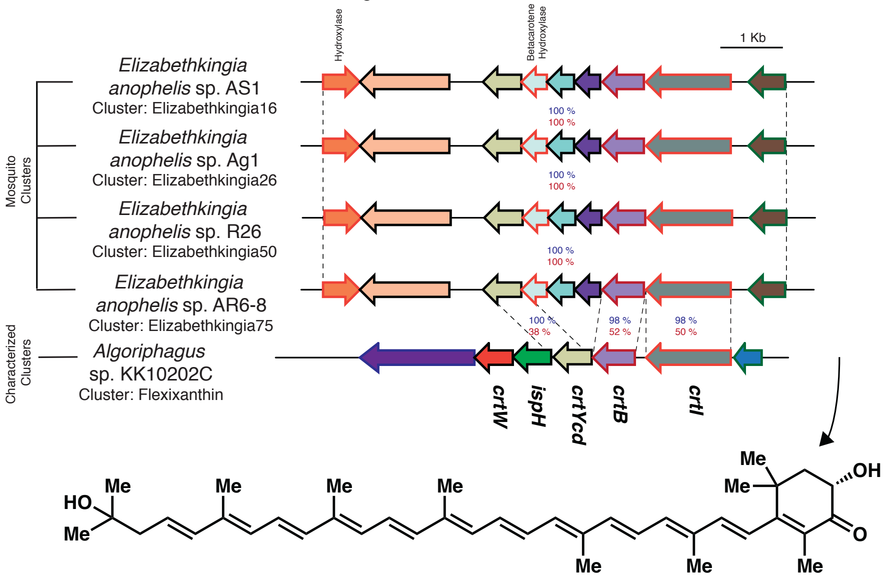


**Flexixanthin**

Kosakonia6:

**Reference**: Sedkova N, Tao L, Rouvière PE, Cheng Q. 2005. Appl Environ Microbiol 71:8141–6.

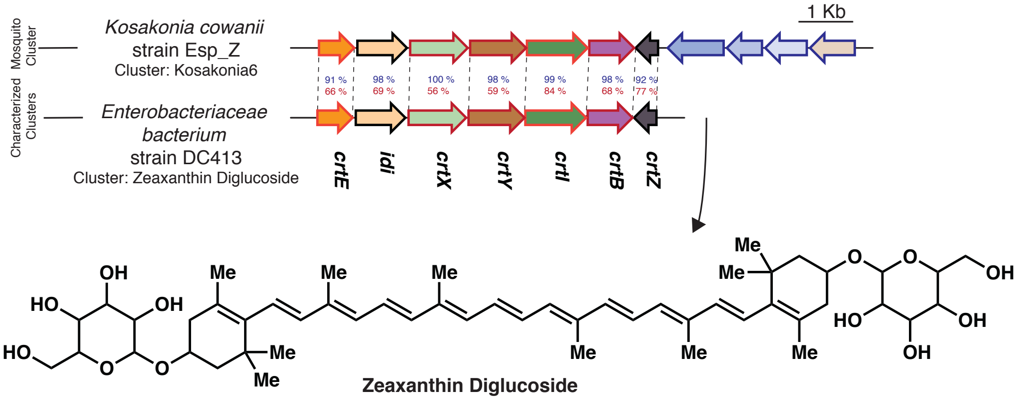


**Zeaxanthin Diglucoside**

Microbacterium41:

**Reference**: Krubasik P, Sandmann G. 2000. Mol Gen Genet MGG 263:423–432.

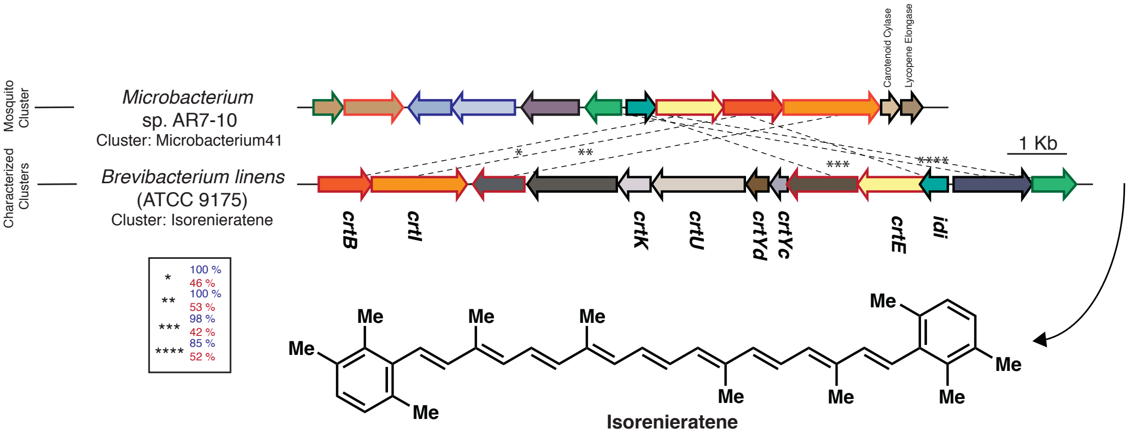


**Isorenieratene**

#### Pantoea6:

**Reference**: Sedkova N, Tao L, Rouvière PE, Cheng Q. 2005. Appl Environ Microbiol 71:8141–6.

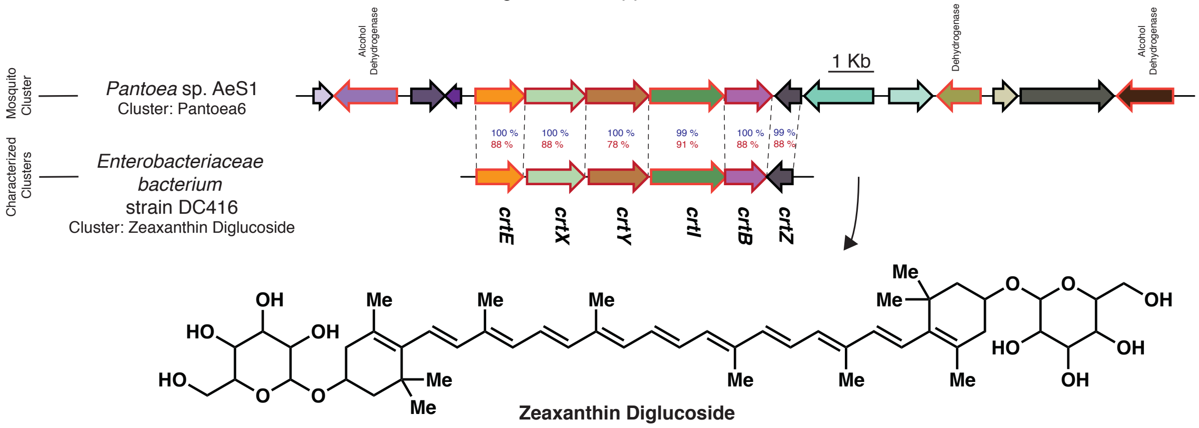


#### Pseudomonas48:

**Reference**: Cimermancic P, Medema MH, Claesen J, Kurita K, Wieland Brown LC, Mavrommatis K, Pati A, Godfrey PA, Koehrsen M, Clardy J, Birren BW, Takano E, Sali A, Linington RG, Fischbach MA. 2014. Cell 158:412–421.

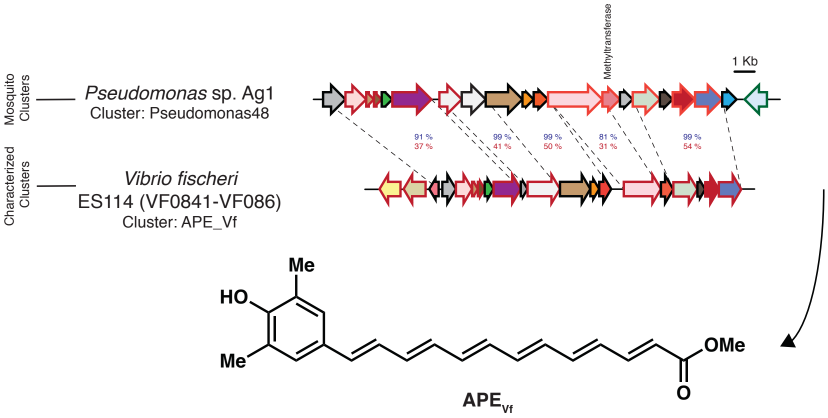


#### Pseudomonas55:

**Reference**: Sedkova N, Tao L, Rouvière PE, Cheng Q. 2005. Appl Environ Microbiol 71:8141–6.

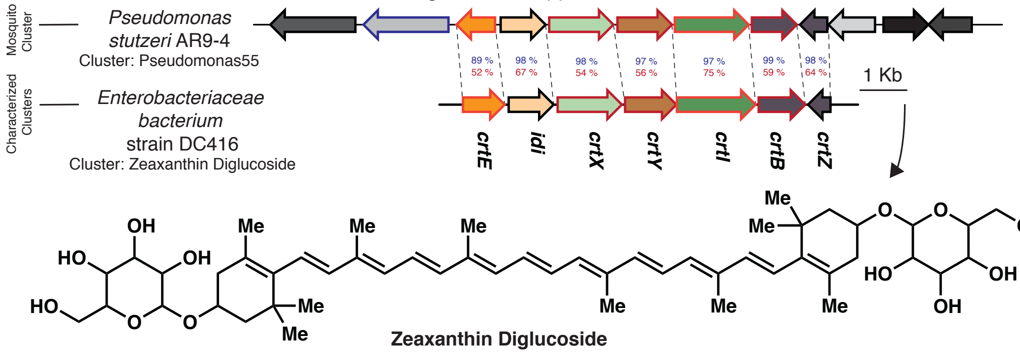

#### Pseudomonas67:

**Reference**: Cimermancic P, Medema MH, Claesen J, Kurita K, Wieland Brown LC, Mavrommatis K, Pati A, Godfrey PA, Koehrsen M, Clardy J, Birren BW, Takano E, Sali A, Linington RG, Fischbach MA. 2014. Cell 158:412–421.

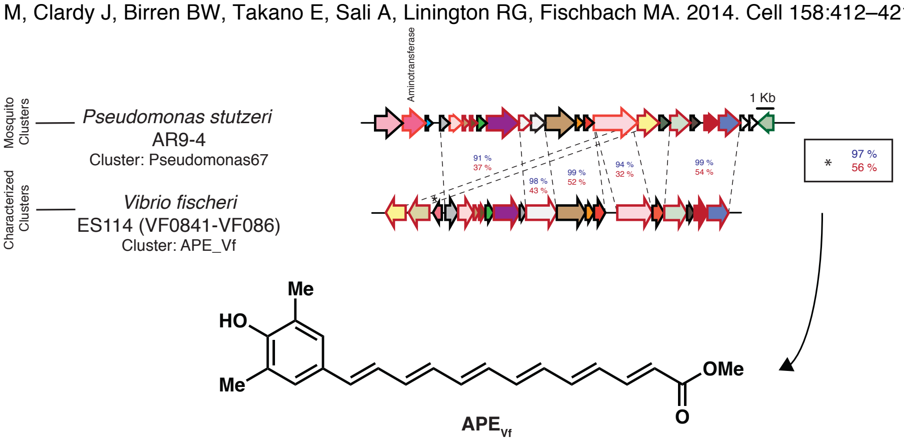


#### Serratia52 & Serratia162:

**Reference**: Cimermancic P, Medema MH, Claesen J, Kurita K, Wieland Brown LC, Mavrommatis K, Pati A, Godfrey PA, Koehrsen M, Clardy J, Birren BW, Takano E, Sali A, Linington RG, Fischbach MA. 2014. Cell 158:412–421.

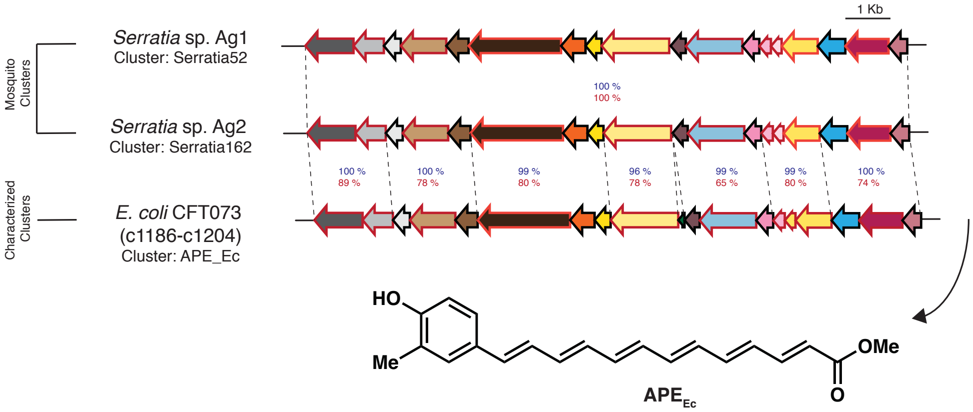


#### Sphingobacterium26:

**Reference**: Tao L, Yao H, Kasai H, Misawa N, Cheng Q. 2006. Mol Genet Genomics 276:79–86.

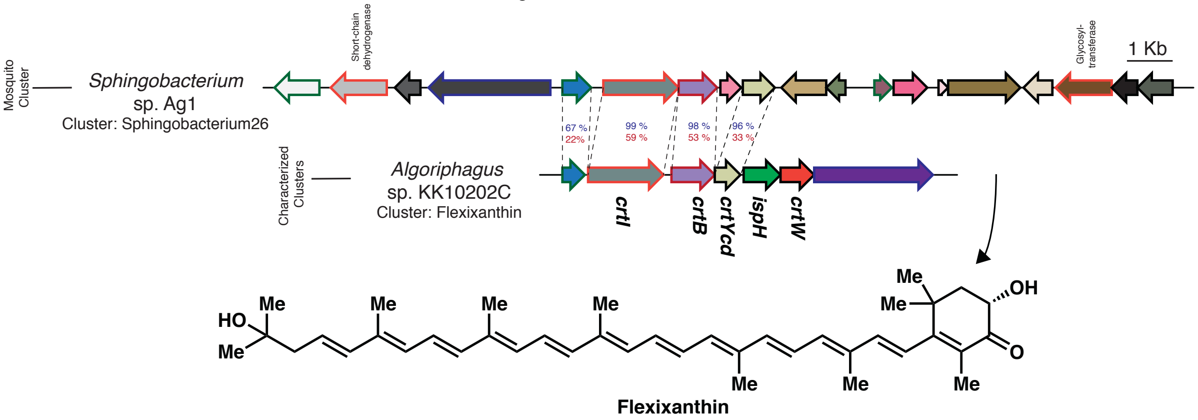

#### Sphingomonas11:

**Reference**: Kim SH, Kim JH, Lee BY, Lee PC. 2014. Appl Microbiol Biotechnol 98:9993–10003.

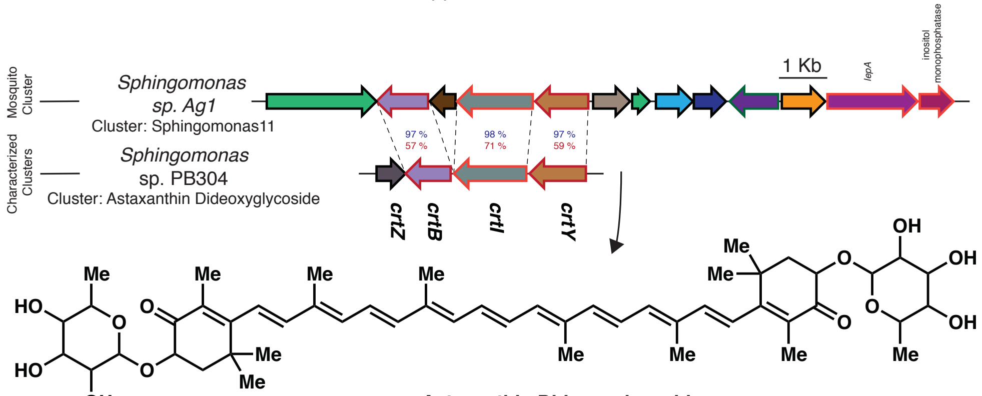


**Astaxanthin Dideoxyglycoside**

#### Staphylococcus5:

**Reference**: Kim SH, Lee PC. 2012. J Biol Chem 287:21575–83.

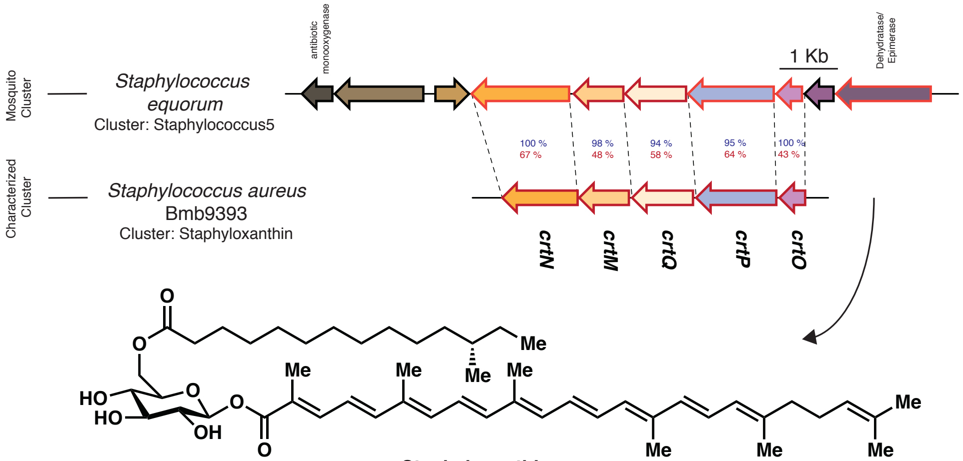


**Staphyloxanthin**

#### Stenotrophomonas16:

**Reference**: Cimermancic P, Medema MH, Claesen J, Kurita K, Wieland Brown LC, Mavrommatis K, Pati A, Godfrey PA, Koehrsen M, Clardy J, Birren BW, Takano E, Sali A, Linington RG, Fischbach MA. 2014. Cell 158:412–421.

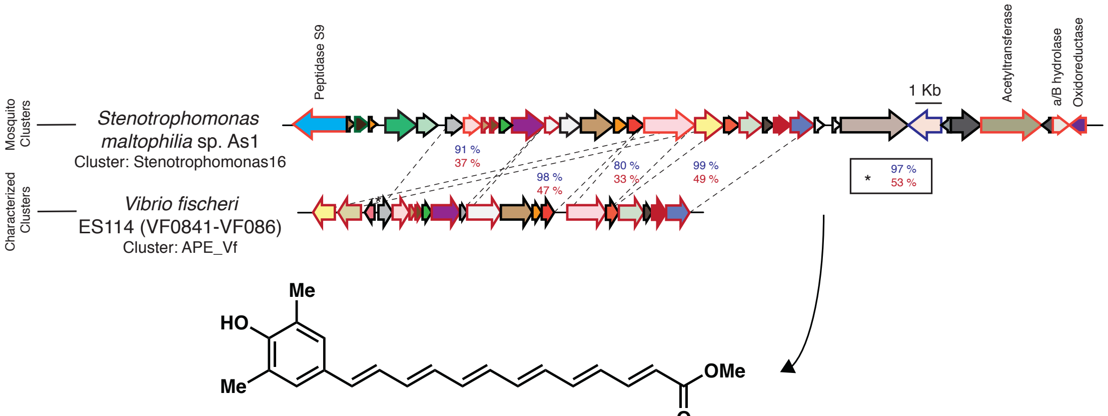


**APE$_{Vf}$**

**Autoinducer:**

<u>Chromobacterium2:</u>

**Reference**: Rekha PD, Young C-C, Arun AB. 2011. 3 Biotech 1:239–245.

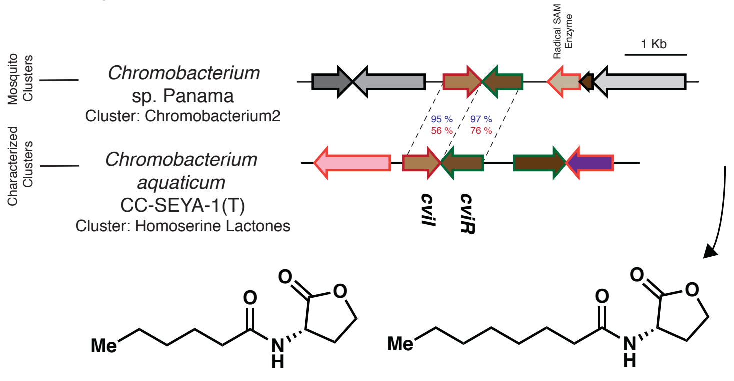


***N*-hexanoyl homoserine lactone** ***N*-octanoyl homoserine lactone**

<u>Enterobacter10:</u>

**Reference**: Lau YY, How KY, Yin W, Chan K. 2018. Microbiologyopen 7:e00610.

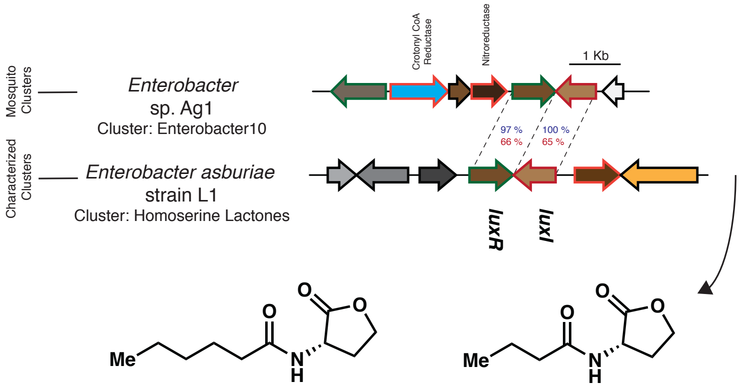


***N*-hexanoyl homoserine lactone** ***N*-butyryl homoserine lactone**

<u>Pantoea15:</u>

**Reference**: Lee J, Zhang L. 2015. Protein Cell 6:26–41.

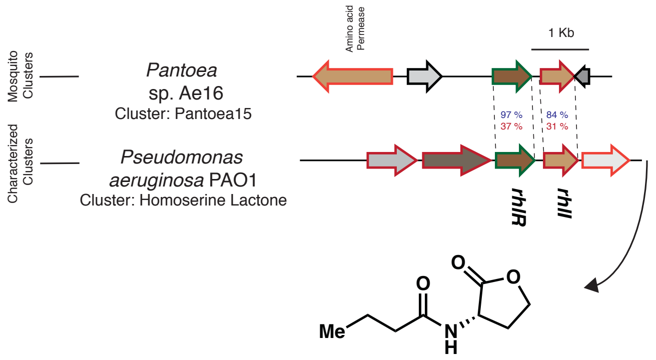


***N*-butyryl homoserine lactone**

#### Pseudomonas19:

**Reference**: Kretsch AM, Morgan GL, Tyrrell J, Mevers E, Vallet-Gély I, Li B. 2018. Org Lett 20:4791–4795.

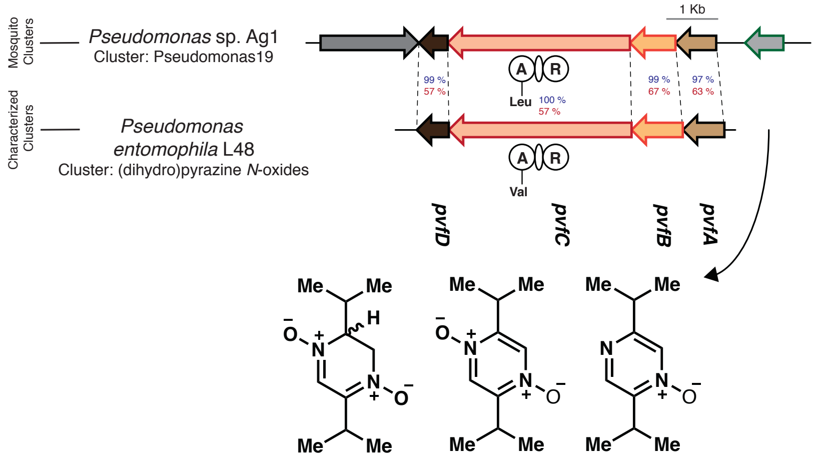


**(dihydro)pyrazine *N*-oxides**

#### Leucobacter5:

**Reference**: Kato J, Funa N, Watanabe H, Ohnishi Y, Horinouchi S. 2007. Proc Natl Acad Sci U S A 104:2378–83.

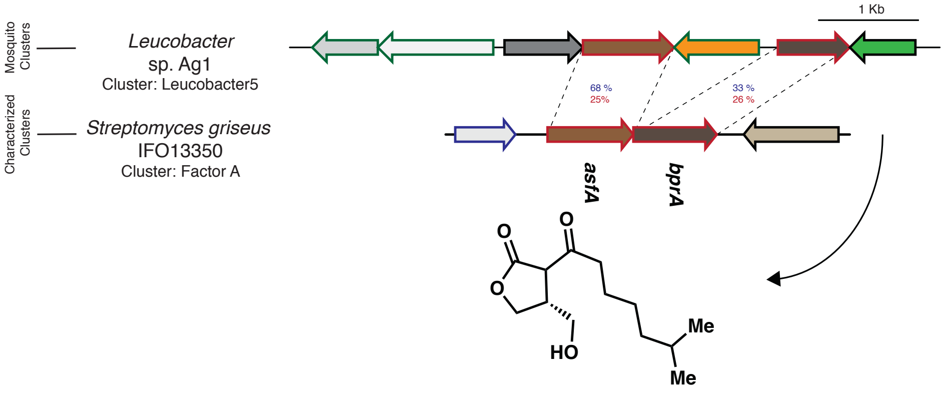


**Factor A**

#### Serratia23, Serratia48 & Serratia180:

**Reference**: Jung BK, Khan AR, Hong S-J, Park G-S, Park Y-J, Park CE, Jeon H-J, Lee S-E, Shin J-H. 2017. J Biotechnol 241:158–162.

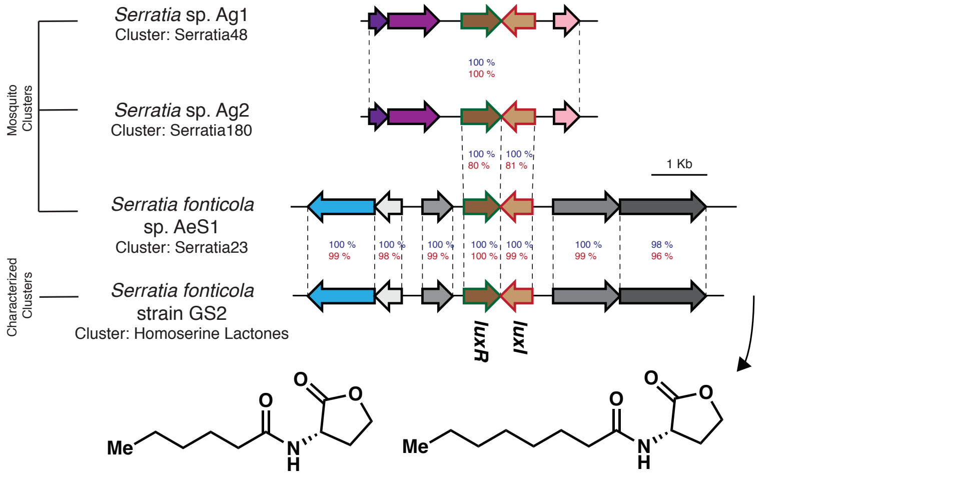


***N*-hexanoyl homoserine lactone** ***N*-octanoyl homoserine lactone**

#### Serratia63, Serratia94 & Serratia125:

**Reference**: Wei J-R, Tsai Y-H, Horng Y-T, Soo P-C, Hsieh S-C, Hsueh P-R, Horng J-T, Williams P, Lai H-C. 2006. J Bacteriol 188:1518–25.

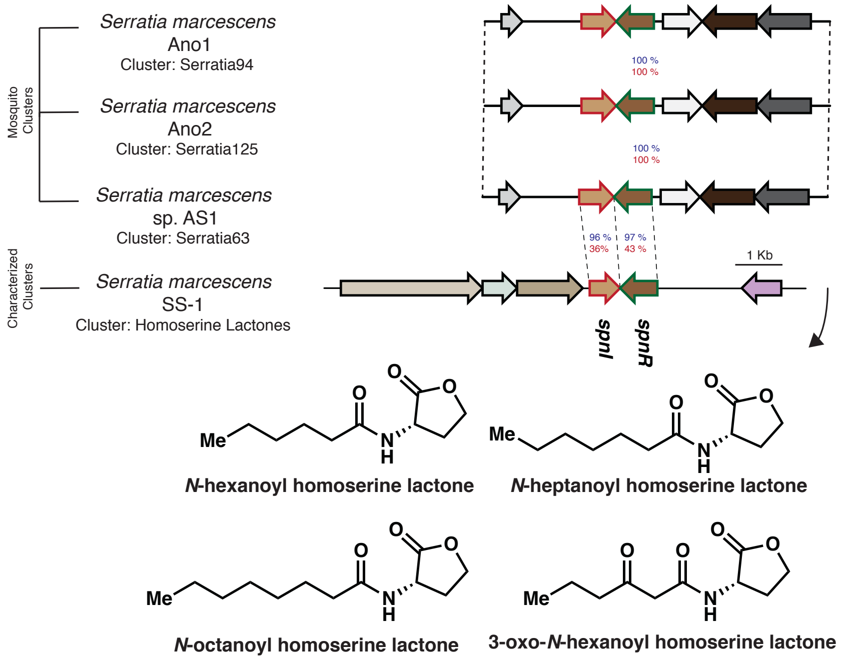


### Antimicrobial:

#### Chromobacterium4:

**Reference**: Saraiva RG, Huitt-Roehl CR, Tripathi A, Cheng Y-Q, Bosch J, Townsend CA, Dimopoulos G. 2018. Sci Rep 8:6176.

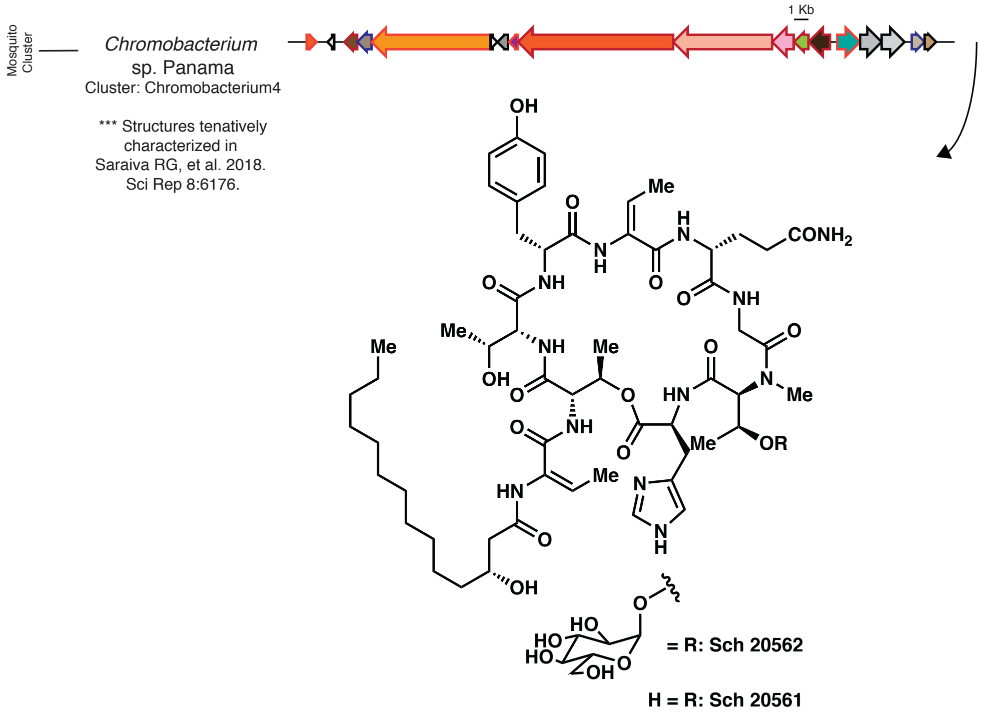

#### Chromobacterium11:

**Reference**: Masschelein J, Clauwers C, Awodi UR, Stalmans K, Vermaelen W, Lescrinier E, Aertsen A, Michiels C, Challis GL, Lavigne R. 2015. Chem Sci 6:923–929.

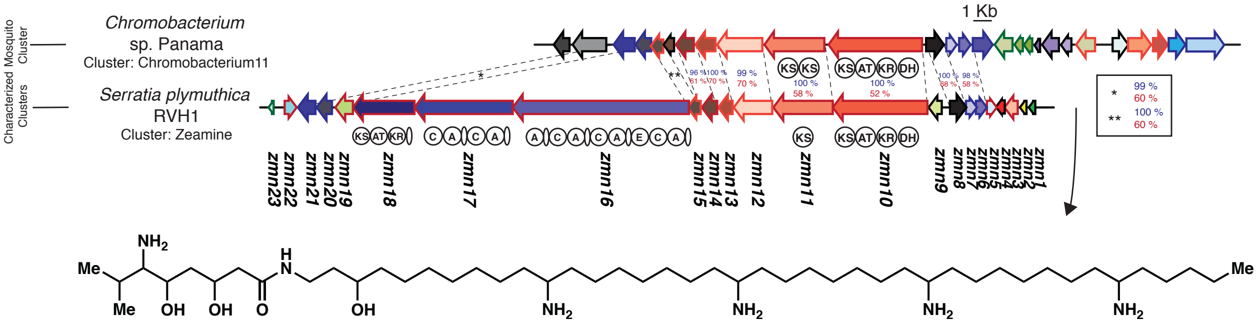


**Zeamine**

#### Pseudomonas1 & Pseudomonas5:

**Reference**: Rokni-Zadeh H, Li W, Sanchez-Rodriguez A, Sinnaeve D, Rozenski J, Martins JC, De Mot R. 2012. Appl Environ Microbiol 78:4826–34.

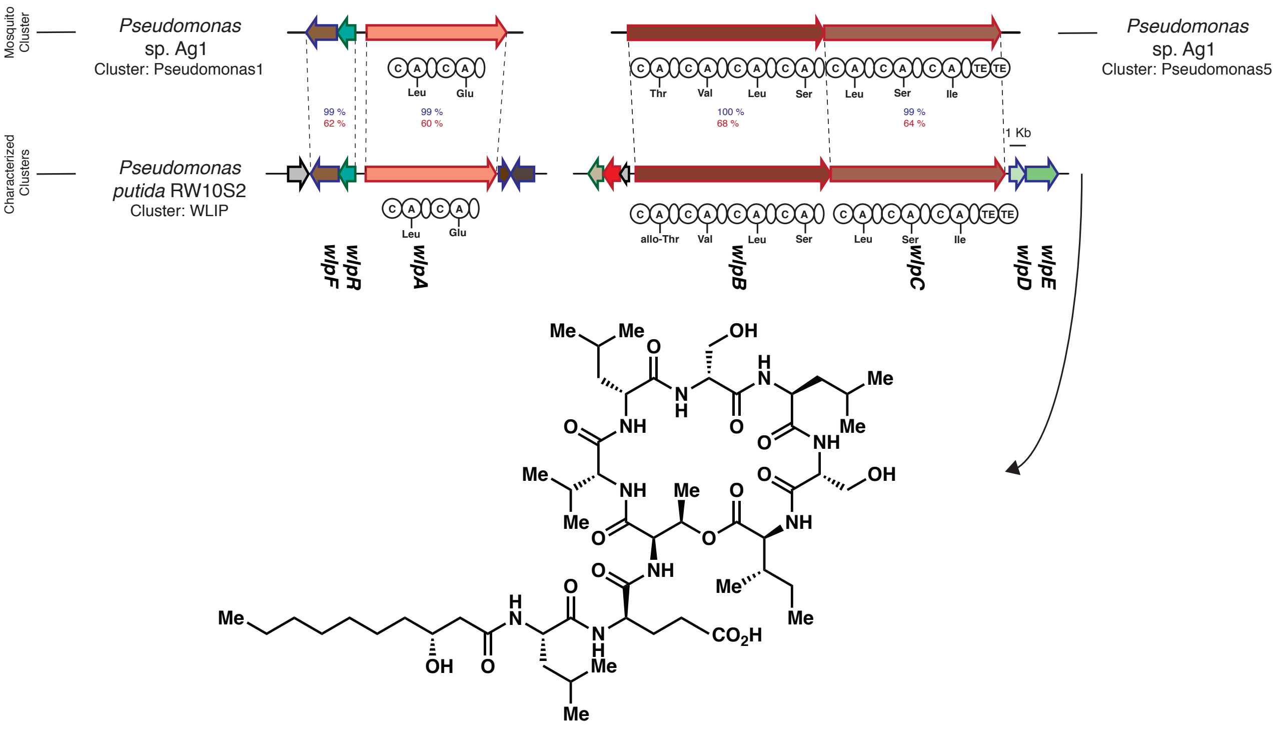


**WLIP**

#### Serratia70, Serratia99, Serratia130 & Serratia151:

**Reference**: Ganley JG, Carr G, Ioerger TR, Sacchettini JC, Clardy J, Derbyshire ER. 2018. ChemBioChem 19:1590–1594.

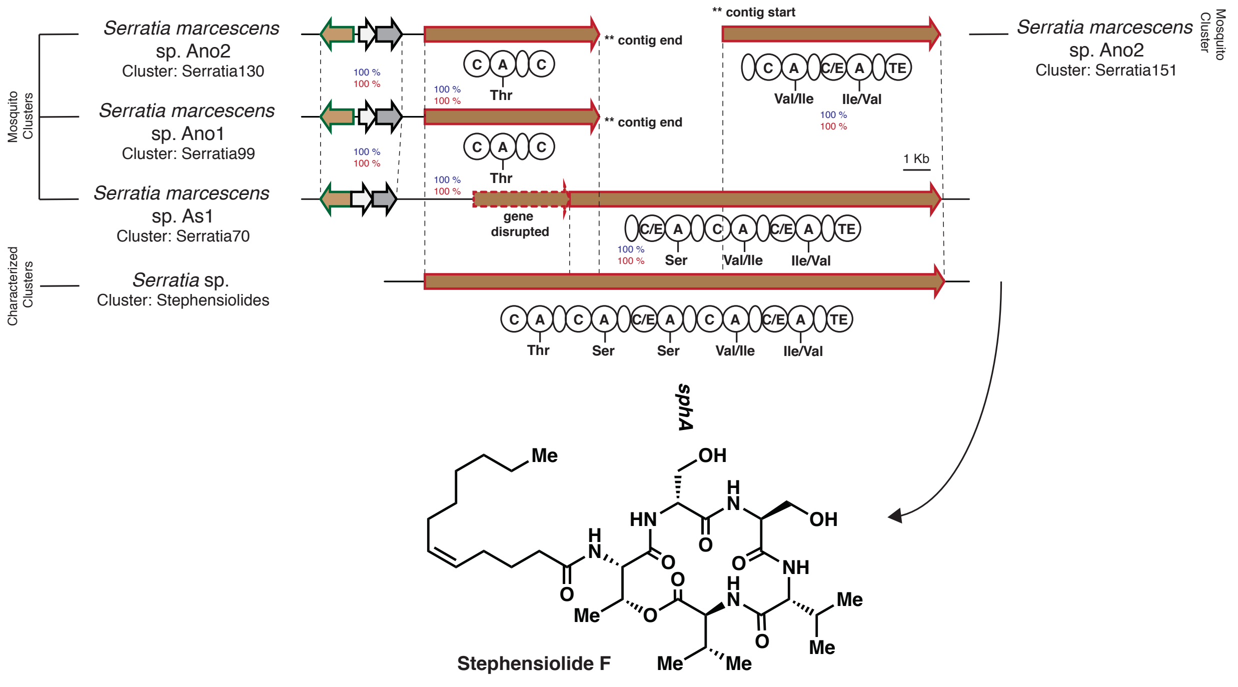


#### Staphylococcus10:

**Reference**: Rodríguez A, Suárez JE, Fernández M, Martínez B. 1999. Microbiology 145:3155–3161

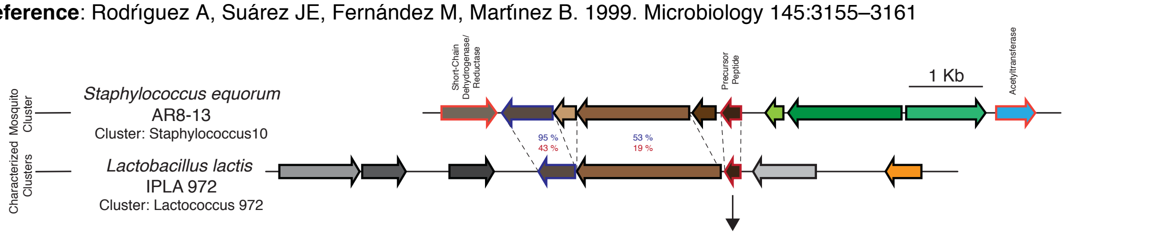


```
Staphylococcus10   MKKTIISVLLYLAIVLGSGTIANAATIYAQG-GIWNYGVGSKYVWSYYSNNYKAHGSTAVGKYKSYSGKREQVSKLEHLPQNHS
                            L + L + T+ +A  I AQ  G W +G G    +S Y +  K H +T V        GK  Q
Lactococcus 972           MKTKSLVLALSAVTLFSAGGIVAQAEGTWQHGYGVSSAYSNYHHGSKTHSATVVNNNTGRQGKDTQRAGVWAKATVGRNLTEKASFYYNFW
```

Signal Peptide

Core Peptide

### Other:

#### Chromobacterium6:

**Reference**: Saraiva RG, Huitt-Roehl CR, Tripathi A, Cheng Y-Q, Bosch J, Townsend CA, Dimopoulos G. 2018. Sci Rep 8:6176.

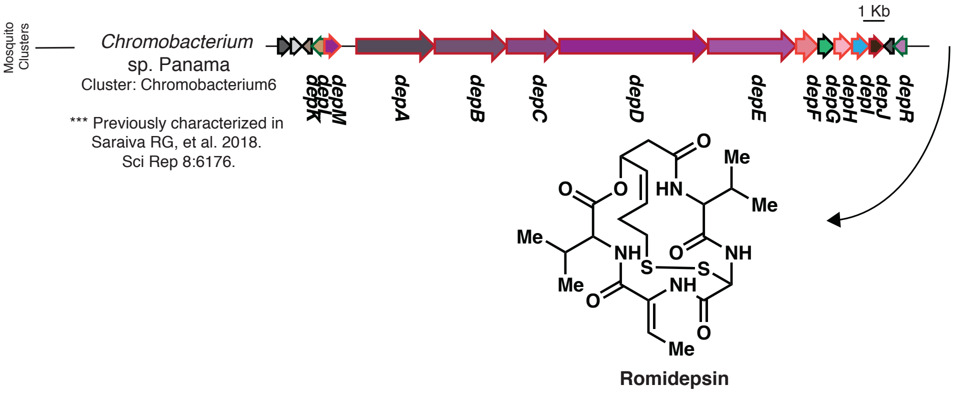

#### Pseudomonas6:

**Reference**: Kuhlmann AU, Bremer E, Le Good JA, Bremer E. 2002. Appl Environ Microbiol 68:772–83.

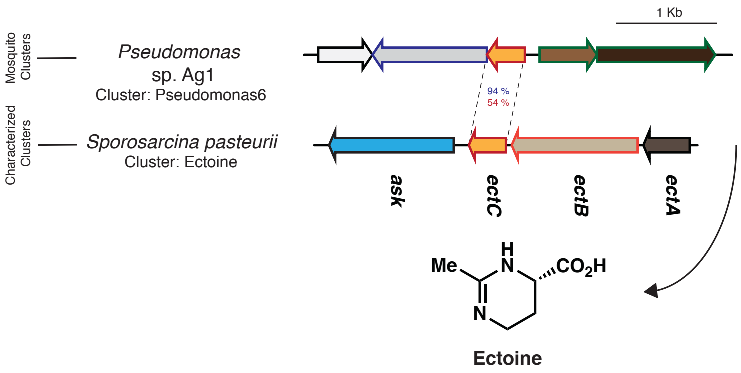


#### Pseudomonas51:

**Reference**: Reshetnikov AS, Khmelenina VN, Trotsenko YA. 2006. Arch Microbiol 184:286–297.

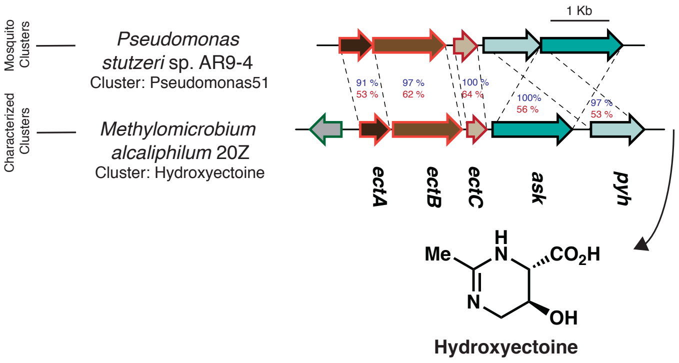


#### Sphingomonas3:

**Reference**: Reshetnikov AS, Khmelenina VN, Trotsenko YA. 2006. Arch Microbiol 184:286–297.

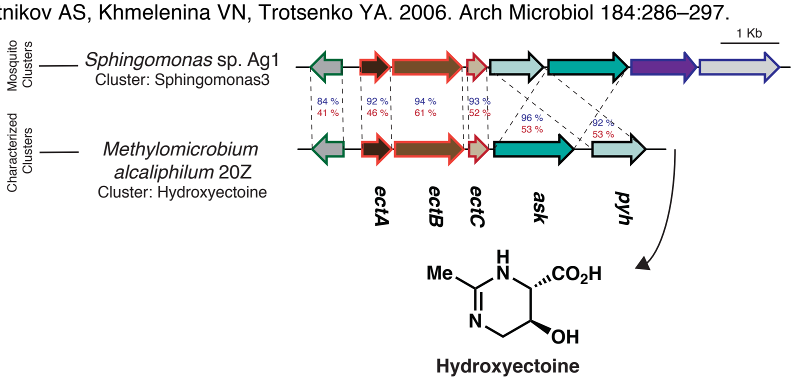

**Resorcinol:**

Leucobacter13:

**Reference**: Funabashi M, Funa N, Horinouchi S. 2008. J Biol Chem 283:13983–91.

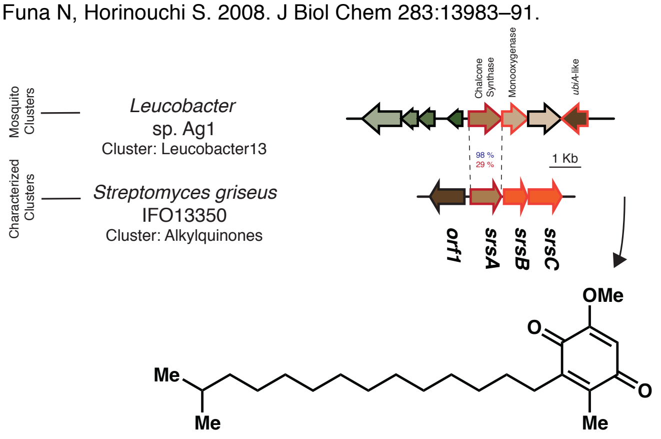


Lysinibacillus26:

**Reference**: Funabashi M, Funa N, Horinouchi S. 2008. J Biol Chem 283:13983–91.

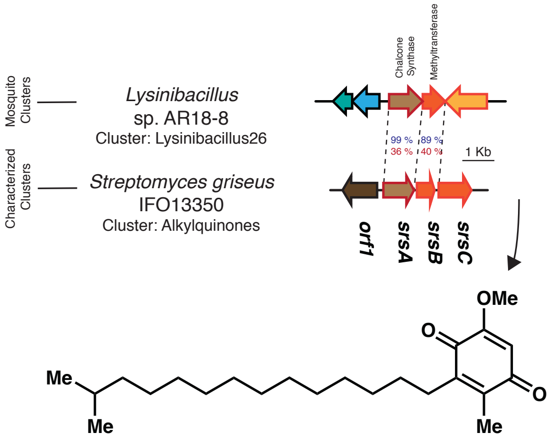


Microbacterium23:

**Reference**: Awakawa T, Fujita N, Hayakawa M, Ohnishi Y, Horinouchi S. 2011. ChemBioChem 12:439–448.

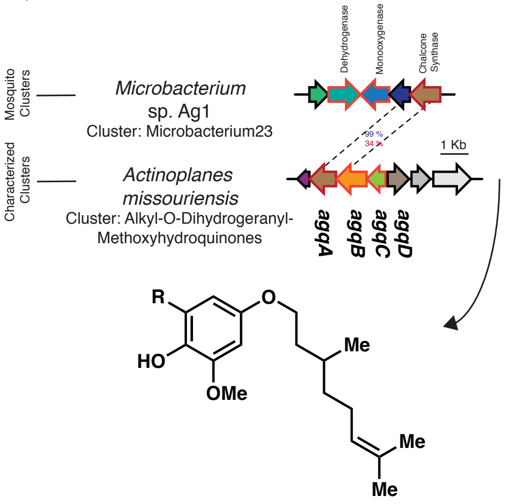

#### Microbacterium32:

**Reference**: Awakawa T, Fujita N, Hayakawa M, Ohnishi Y, Horinouchi S. 2011. ChemBioChem 12:439–448.

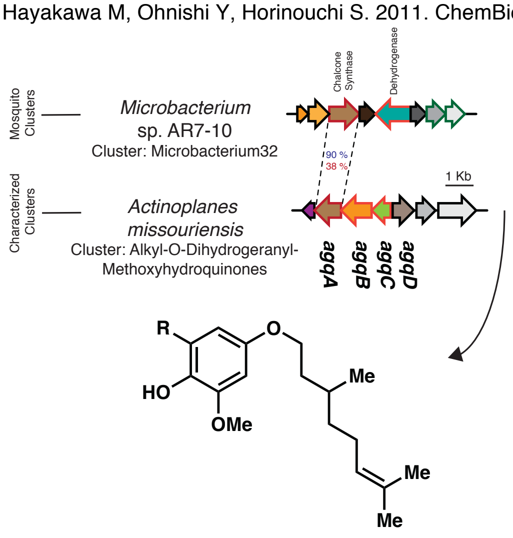


#### Sphingomonas7:

**Reference**: Awakawa T, Fujita N, Hayakawa M, Ohnishi Y, Horinouchi S. 2011. ChemBioChem 12:439–448.

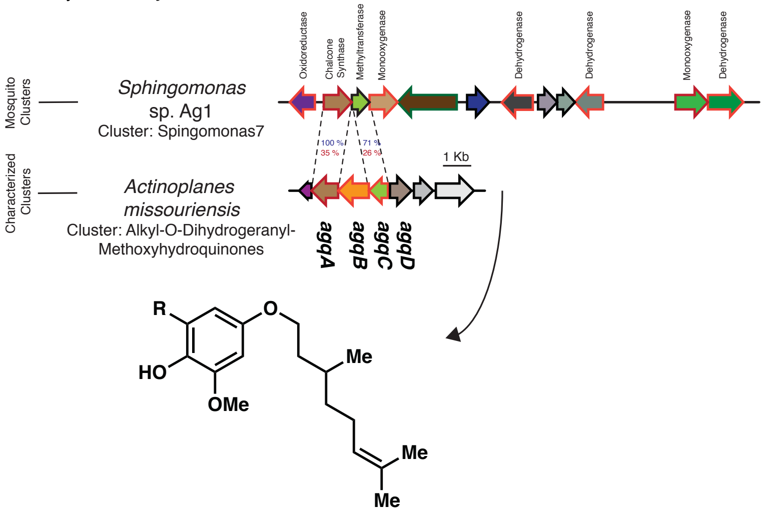


### Lipid/Sterol:

#### Asaia15:

**Reference**: Poralla K, Muth G, HÃ¤rtner T. 2000. FEMS Microbiol Lett 189:93–95.

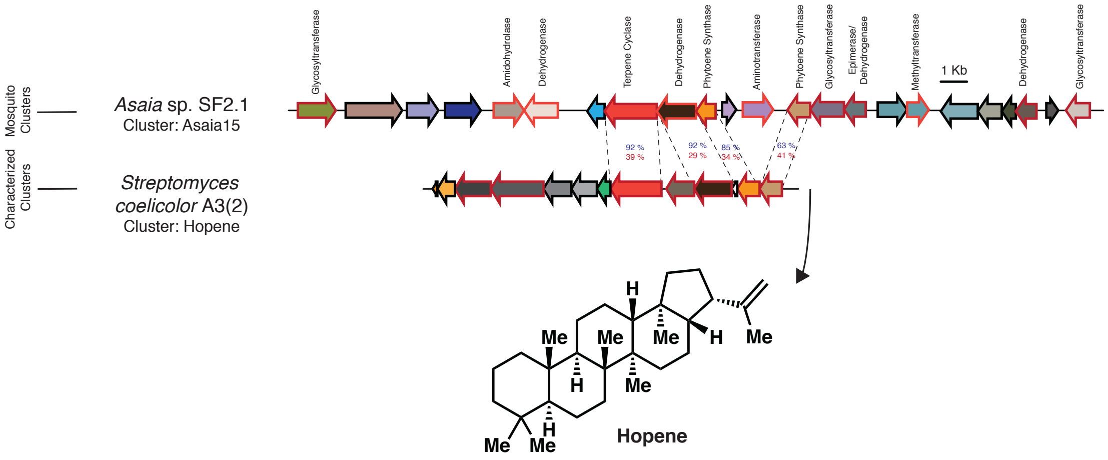

Chromobacterium7:

**Reference:** Pan J-J, Solbiati JO, Ramamoorthy G, Hillerich BS, Seidel RD, Cronan JE, Almo SC, Poulter CD. 2015. ACS Cent Sci 1:77–82.

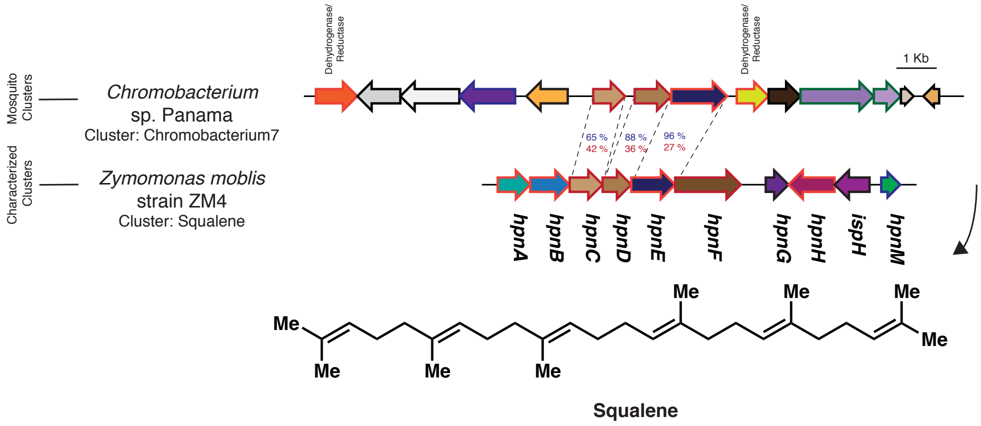
